# Na^+^/Ca^2+^ exchanger triggers transient disruption of axon initial segments in hippocampal granule cells after brief ischemia

**DOI:** 10.1101/2024.12.13.628404

**Authors:** Carol Serra, Emma Martínez-Alonso, María I Ayuso, Ragunathan Padmashri, Christian Henneberger, Dmitri A. Rusakov, Alberto Alcázar, Ricardo Scott

**Affiliations:** Instituto de Neurociencias UMH-CSIC, Alicante, SPAIN; Hospital Ramón y Cajal, IRYCIS, Madrid, SPAIN; UCL Institute of Neurology, University College London, London, UK; Institute of Cellular Neurosciences, University of Bonn, Bonn, Germany; German Center for Neurodegenerative Diseases (DZNE), Bonn, Germany

**Keywords:** axon initial segment, Na^+^/Ca^2+^ exchanger, ischemia, calpain

## Abstract

The axon initial segment (AIS) is the site of action potential initiation in many cell types. The large density of Na^+^ channels and the small intraluminal volume of the AIS underlie the largest spike-dependent Na^+^ raise along the neuron. Our Na^+^ and Ca^2+^ imaging experiments in dentate granule cells reveal that the Na^+^/Ca^2+^ exchanger contributes to fast Na^+^ clearance at the expense of Ca^2+^ entry specifically at the AIS during short bursts of action potentials. The AIS is thought to be irreversibly disrupted during brain ischemia by the Ca^2+^- dependent protease calpain, as an early step leading to neuronal death. We find here that brief transitory ischemia produces a similar calpain-dependent disruption of the AIS in the dentate gyrus. However, this is not an irreversible process: a few days following the brief ischemic stress, intact initial segments re-appear, and calpain activity in the dentate gyrus returns to normal. Moreover, pharmacological blockade of Ca^2+^ entry through the Na^+^/Ca^2+^ exchanger prevents AIS disruption *in vivo* and in slice preparations. Our results unveil a highly dynamic anoxia driven disruption-reconstruction of AIS, which is mediated by a previously unnoticed Ca^2+^ entry through the Na^+^/Ca^2+^ exchanger in the axon initial segment of hippocampal granule cells.

---

The axon initial segment (AIS) is the preferred site for action potential initiation in most neurons (Araki & Otani, 1955; Coombs *et al*., 1957). Spike-dependent intracellular Na^+^ changes at the AIS are the most prominent Na^+^ increases across the entire neuron (Kole *et al*., 2008; Fleidervish *et al*., 2010). At the same time, spiking sharply raises Ca^2+^ concentration at the AIS (Bender & Trussell, 2009). The Na^+^/Ca^2+^ interplay is however poorly understood at this crucial region of the neuron. In presynaptic axonal compartments, large voltage-dependent changes of free intracellular Ca^2+^ are typically curtailed by proteins like cabindin (Blatow *et al*., 2003; Scott & Rusakov, 2006). A similar machinery is expected to act at the AIS, as these Ca^2+^ buffers are highly diffusible (Blatow *et al*., 2003). However, the mechanisms underlying removal of Na^+^ from the axonal compartments remain to be established. The Na^+^/Ca^2+^ exchanger (NCX) is a fast transporter that exchanges three Na^+^ ions for one Ca^2+^ ion across the plasma membrane. The exchange mechanism is bidirectional, it depends on the ionic concentrations at both sides of the membrane, and acts in a voltage-dependent manner and without direct energy consumption (Blaustein & Lederer, 1999). In normal conditions, the NCX contributes to remove excessive Ca^2+^ from the cytoplasm (forward mode), especially at the postsynaptic site but also in presynaptic terminals (Roome *et al*., 2013). With repetitive stimulation the NCX can reverse its mode of action at presynaptic terminals and even contribute to transmitter release (Minami *et al*., 2007; Roome *et al*., 2013). Despite the intense fluxes of Na^+^ and Ca^2+^ at the AIS, a local involvement of NCX has remained unknown.

The AIS is far from being a static region: it changes its structure and function depending on cellular and network activity (Grubb & Burrone, 2010; Gutzmann *et al*., 2014). Changes in AIS have been related to neurological disease such as epilepsy and stroke (Wimmer *et al*., 2010; Yoshimura & Rasband, 2014). Transient ischemia produces irreversible proteolysis of several proteins forming the AIS leading to neuronal death (Schafer *et al*., 2009).

In myelinated axons, whose nodes of Ranvier resemble the AIS, NCX reverse mode has an important role in axon degeneration (Stys *et al*., 1991). Under metabolic stress axons progressively lose control of their membrane potential, becoming depolarised and producing a persistent Na^+^ entry through low-threshold/persistent-current Nav1.6 sodium channels. Then, the NCX tries to re-establish Na^+^ at the expense of introducing Ca^2+^ into the axons, leading to axonal degeneration and cell death (Stys *et al*., 1992; Barsukova *et al*., 2012; Bei & Smith, 2012; Zhang & David, 2016).

Here we explore hippocampal granule cells (CGs) and find a prominent spike-dependent Ca^2+^ entry mediated by the NCX, in normal conditions, specifically at the AIS. Under anoxia, global Ca^2+^ entry is also facilitated by NCX whereas its pharmacological blockade reduces AIS disintegration under ischemic stress. Our data thus suggest that NCX plays a major role in the ionic homeostasis at the AIS and that its activity underlies rapid disassembly of the AIS during ischemia.

## RESULTS

### Na^+^/Ca^2+^ exchanger at the AIS

To test if NCX plays a role in Na^+^/Ca^2+^ interplay at the AIS during spiking activity, we imaged Na^+^ and Ca^2+^ entry induced by short trains of APs and tested the effect of KB-R7943, a potent inhibitor of the NCX reverse mode (NCX_rev_; Na^+^ efflux mode). Somatic stimulation of GCs induces prominent spike-dependent Na^+^ elevations (imaged with SBFI, Methods) at the AIS (Kole *et al*., 2008; Scott *et al*., 2014) (Fig. 1A&C). Subthreshold depolarization also induces large Na^+^ transients in this region (Fig.1B). The specific NCX_rev_ blocker KB-R7943 (2μM) increased spike-induced Na^+^ transients by 53±11% (n = 8, p < 0.01; Fig.1A&C-D). In contrast, in cells loaded with the Ca^2+^ sensitive dye Fluo-4, the drug *reduced* Ca^2+^ transients induced by identical stimuli by 23±4% (n = 6, p<0.01; Fig.1E-G). Moreover, with a high internal Na^+^ concentration ([Na^+^]_I_=28mM), the reduction of Ca^2+^ transients by KB-R7943 was more prominent (40±5%, n = 10, p < 0.01; Fig. 1F-G). These series of effects (Fig.1D&G) are fully consistent with the recruitment of NCX_rev_ (Na^+^- extrusion/Ca^2+^-entry) by short trains of APs.

**Figure 1.**
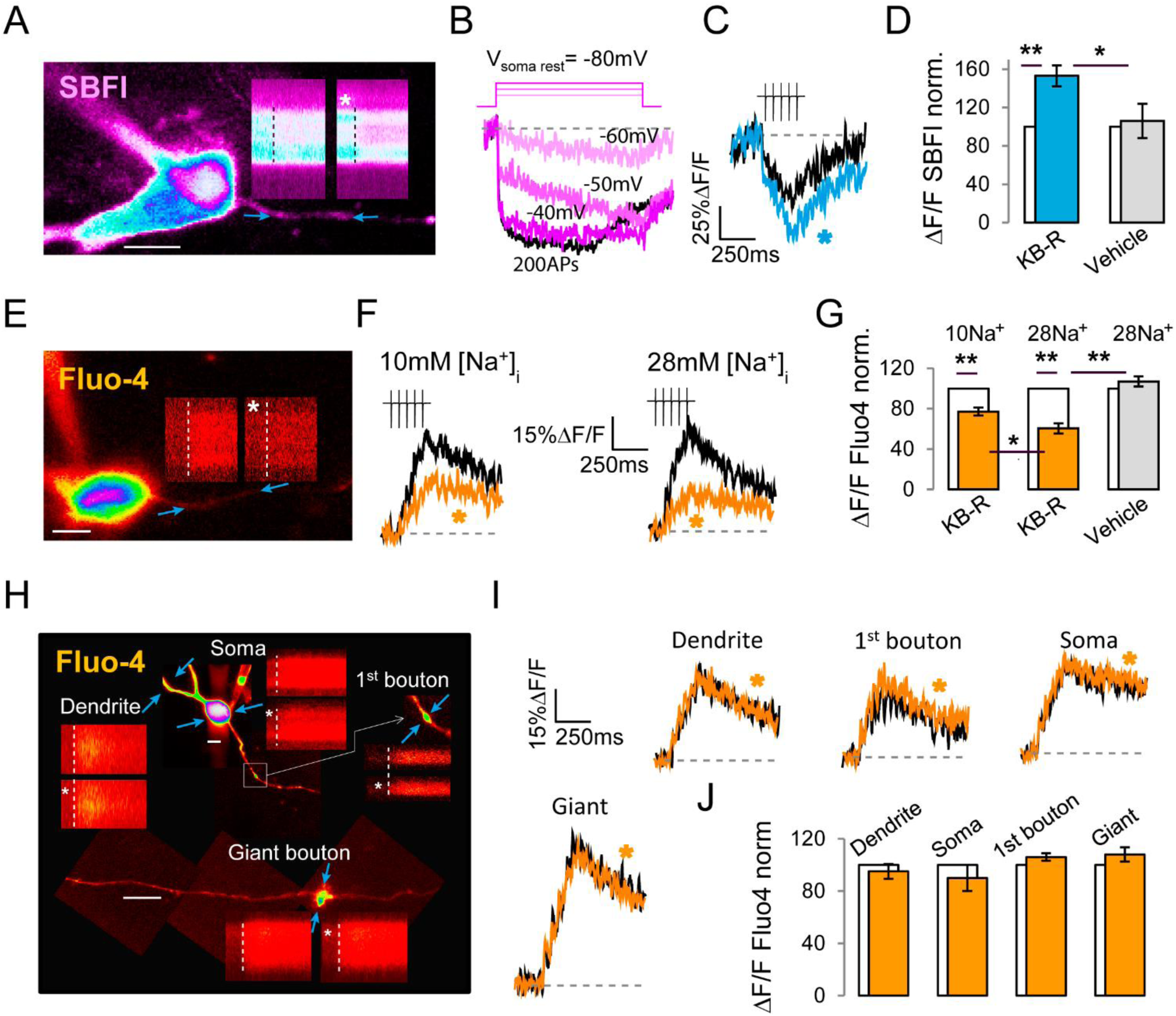
Brief bursts of spikes induce Na^+^ efflux in exchange of Ca^2+^ entry at the AIS through the NCX in hippocampal GCs. (**A**) Two-photon image of a GC loaded with SBFI (1mM) through the patch pipette. Insets are linescans along the AIS (arrows) revealing Na^+^-dependent fluorescent transients induced by 5 somatic pulses at 20Hz in voltage clamp (starting at the dotted line). The asterisk indicates bath application of 2μM KB-R7943, which produces an increase in Na^+^ influx. (**B**) Representative SBFI signals in response to different amplitudes of voltage somatic pulses (as voltage protocol) and a train of high frequency APs (black trace). (**C**) Traces corresponding to linescans shown in (A), and the effect of KB-R7943 (blue, asterisk). (**D**) Average effects of KB-R7943 (n=8), and vehicle (n=6) with respect to control (before bath application). (**E**) GC loaded with Fluo-4 (200μM) and Alexa Fluor-598 (40μM). Insets show Ca^2+^-dependent fluorescent transients in response to the same stimulating protocol as in (C), and the effect of KB-R7943. (**F**) Traces corresponding to transients in 10mM (*left*) and 28mM internal [Na^+^] (*right*). (**G**) Summary of the effect of KB-R7943 for experiments in 10mM Na^+^_i_ (n=6), 28mM Na^+^_i_ (n=10), and vehicle (n=8). (**H**) Examples of linescans from different regions of GCs, as indicated, and the effect of KB-R7943 (asterisks). (**I**) Traces corresponding to Ca^2+^ transients from those regions, as indicated. Asterisks indicate the presence of KB-R7943. (**J**) Summary of the effect of KB-R7943 on Ca^2+^ transients from each of the regions (n=6 per region). Data are mean ± SEM, *p<0.05 **p<0.01. Scale bars in pictures are 5 μm.

Next, we tested whether other regions of the neuron displayed a similar activity of NCX with the same stimulation protocol. We recorded Ca^2+^ transients through-out the neuron before and after bath application of KB-R7943 (2 μM) in the presence of high [Na^+^] in the pipette to enhance NCX reverse activity. However, Ca^2+^ transients at the soma, dendrites, proximal axonal boutons and giant boutons in CA3 area were not affected by the drug. This indicates that activation of NCX_rev_ by short bursts is a specific phenomenon of the AIS; and further confirms the specificity KB-R7943 effects on Ca^2+^ signals at the AIS.

As the patch pipette dialysis could potentially affect the endogenous ion dynamics at the AIS, we designed an experiment to evaluate the effect of KB-R7943 on Ca^2+^ transients in axons left naturally to regulate their Na^+^ concentration (Supplementary Fig.1). We thus patched CGs and loaded them with 400μM Fluo-4 and 80μM Alexa Fluor-594 for 2-3 minutes. We then pulled-out the pipette to reseal the cell and waited for 30 min at 34°C to let the Na^+^_i_ concentration reestablish. Next, we re-patched the cell in cell-attached configuration and stimulated the axon with 5 pulses at 20 Hz, by means of a bipolar electrode inserted in the stratum lucidum (∼500 μm far away from the soma). The stimulation protocol induced similar Ca^2+^ transients to those observed in whole-cell mode, and KB-R7943 produced a ∼20% reduction of the transients amplitude. The comparison of these effects with the results shown in Fig.1G, suggests that the “natural” intracellular Na^+^ concentration in slices is ∼10mM and that the NCX can operate in the reverse-mode in the AIS in cells left to regulate their membrane potential by themselves.

Next, we tested the effect of additional NCX_rev_ blockers (Supplementary Fig. 2). First, we cross-tested the pharmacological effects of KB-R7943 with the peptide XIP, a blocker of NCX_rev_, which works predominantly from inside the cells. XIP (1μM) was included in a 28mM Na^+^ internal solution and loaded through the patch pipette into the cells. We monitored AIS-Ca^2+^ influx after 3 minutes from breaking-in, well before the peptide XIP could reach a maximal concentration at the AIS. After 15 minutes, we resumed Ca^2+^ transient recording. Spike-induced Ca^2+^ transients were reduced by 30±11% (n = 6, p<0.05) in this second time window (high XIP) with respect to the first (low XIP). However, in the presence of high XIP (15 min after breaking-in), Ca^2+^ transients were not altered when KB-R7943 was applied to the bath. The occlusion of the effect of KB-R7943 by XIP is, once again, consistent with a specific effect of KB-R7943 on the NCX_rev_.

We also tested the effect of SN-6, a blocker more specific for NCX1 (Iwamoto *et al*., 2004). This drug significantly reduced Ca^2+^ entry by 23±4% (n = 6, p<0.05), as compared to the 40% reduction with KB-R7943 (Supplemental Fig.2). As KB-R7943 preferentially blocks the NCX3 subtype of the transporter (Iwamoto & Shigekawa, 1998), this result, besides confirming a spike-dependent activation of NCX_rev_, it suggests that the major subunit composition of the AIS exchanger could be NCX3.

**Figure 2.**
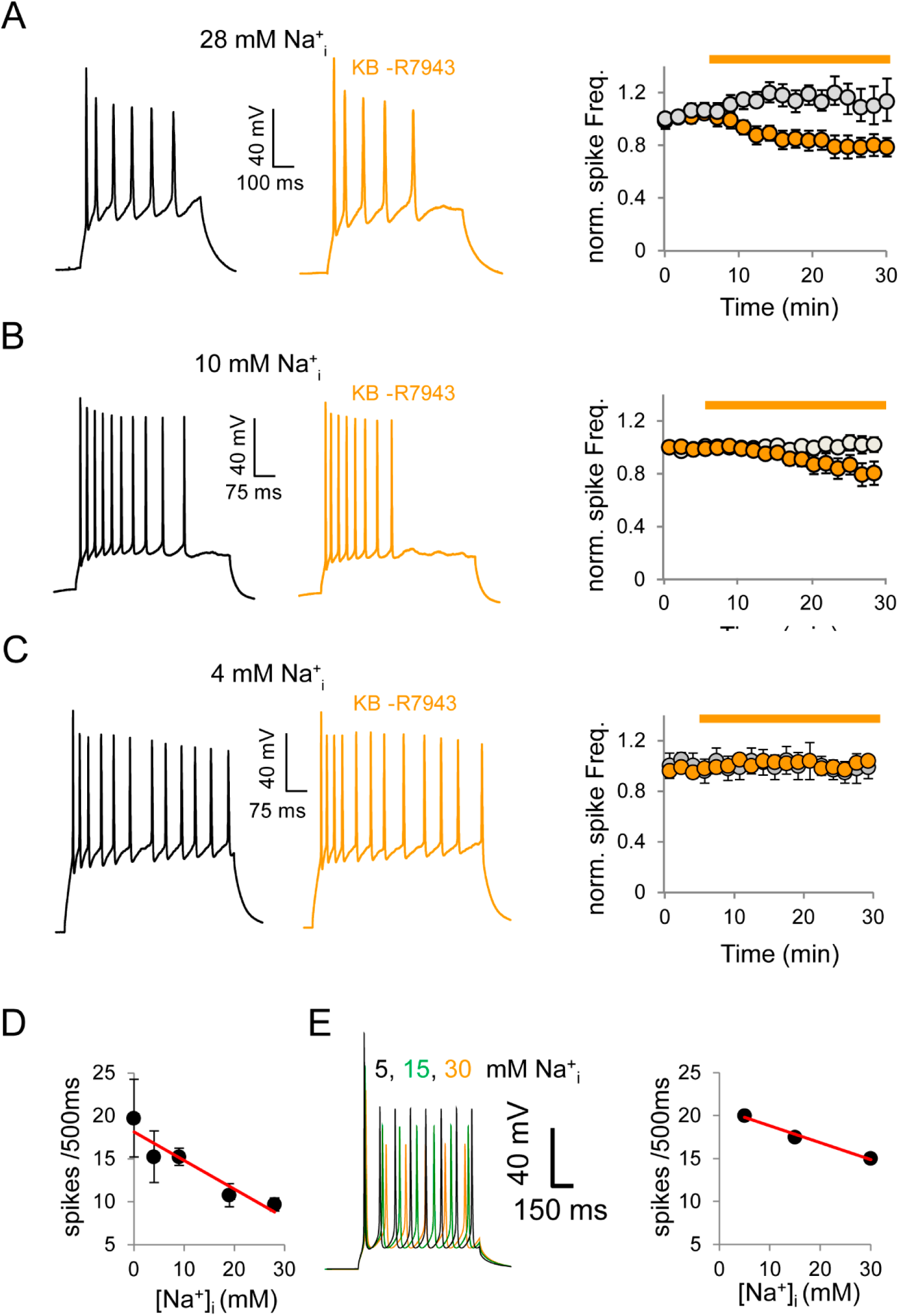
NCX_rev_ increases the firing frequency of granule cells by augmenting the Na^+^ driving force depending on internal [Na^+^]. (**A**) Firing sweeps in response to somatic depolarization in cells loaded with high [Na^+^]_i_ (28mM) before and after (orange) 10min of bath-applied KB-R7943 (2 μM). *Right:* average effect of KB-R7943 on spike frequency. Grey dots correspond to vehicle perfusion (data are normalized; n=14 per condition). (**B**) Effect of KB-R7943 or vehicle on spike frequency in lower [Na^+^]_i_ (10mM Na^+^_i_; n=9 per condition). (**C**) Effect of KB-R7943 or vehicle on spike frequency in 4mM Na^+^_i_ (n=7 per condition). (**D**) GC firing frequency at different internal [Na^+^], ranging from 0 to 28 mM. (**E**) *Left:* GC firing in different [Na^+^]_i_ in a GC model (Schmidt-Hieber *et al*., 2008; Scott *et al*., 2014). *Right:* Relation between [Na^+^]_i_ and spike frequency. Linear fits in red. Data are average ±SEM. *p<0.05,**p<0.01

### NCX mediated changes in spike frequency

We reasoned that the observed Na^+^ and Ca^2+^ fluxes mediated by NCX at the AP initiation site could have an impact on spike firing. To test the hypothesis, we recorded trains of APs in response to somatic depolarization in cells loaded with high [Na^+^]_i_ (28mM). During KB-R7943 bath perfusion, spike frequency decreased by 26±9% with respect to the vehicle effect (Fig. 2A). As expected from a blockade of NCX_rev_, the effect of KB-R7943 was less prominent in the presence of lower [Na^+^]_i_ (10mM Na^+^_i_), and absent in 4mM Na^+^_i_ (Fig. 2B&C). How does AIS-NCX affect spike frequency? The total amount of Na^+^ entering the AIS during a train of 5 APs can reach 2-5 mM (Shu *et al*., 2006; Scott *et al*., 2014). According to our Na^+^ imaging data, NCX would be responsible for the extrusion of 40% of that Na^+^ entry (Fig.1 main text; increase of 1-2 mM). To test if the effect of NCX blockade on spike frequency could be mediated by local E_Na_ changes, we compared GC firing frequency at different [Na^+^]_i_, ranging from 0 to 28 mM (Fig. 2D). Indeed, firing frequency was reduced with increasing [Na^+^]_i_ (slope -0,33± 0,07, p<0.015). In addition, in a GC model (Schmidt-Hieber *et al*., ^2^00^8^; Scott *et al*., 2014) we obtained a similar relationship between [Na^+^]_i_ and spike frequency (Fig. 2E)

Although in whole-cell mode we only stimulate directly one GC in the slice, bath-applied KB-R7943 might affect the network activity, thus altering firing properties of the recorded GC. To test for this possibility, we repeated the experiments from Fig. 2 with KB-R7943 inside the pipette and with a high concentration of Na^+^ (28 mM). KB-R7943 was applied to the bath, but in these conditions the number of spikes per pulse was not affected, suggesting that NCX-mediated effects on spike frequency are cell specific (Suppl. Fig 3). In summary, we unveil a potent contribution of NCX_rev_ to Na^+^/Ca^2+^ fluxes, spatially restricted to the AIS, with an effect on cell firing, suggesting that AIS NCX contributes to preserve normal firing properties.

### AIS disruption upon oxygen deprivation

Intriguingly, a recent study demonstrated that focal ischemia induces irreversible Ca^2+^/calpain-dependent dismantling of the AIS in the cortex, and that this process is an early stage to neuronal death (Schafer *et al*., 2009). We therefore hypothesized that AIS-NCX_rev_ could be related to AIS disintegration and, ultimately, cell death under ischemic stress.

To enable single-cell probing, we first established that AIS disruption did robustly occur in hippocampal acute slices under oxygen and glucose deprivation (OGD, Methods; Fig.3). Immunostaining in *acute slices* displayed enough amount of preserved AIS for evaluation, even when they suffer an unavoidable deterioration during the slicing procedure (see methods for AIS counting protocol). One hour after OGD, AIS immunostaining for ankyring-G, pIKBα and Na^+^ channels, nearly disappeared specifically from the GC layer, consistent with the disruption, conformational change or removal of these proteins. In contrast, AISs from CA3 and CA1 regions remained intact. Intriguingly, KB-R7943 prevented AIS disruption in GCs (Fig.3). Additionally, a milder and longer OGD (Methods) produced similar results: it disrupted GC-AIS but did not affect CA1 and CA3 AISs; and GC-AIS were also protected by KB-R7943 (Suppl. Fig. 4).

**Figure 3.**
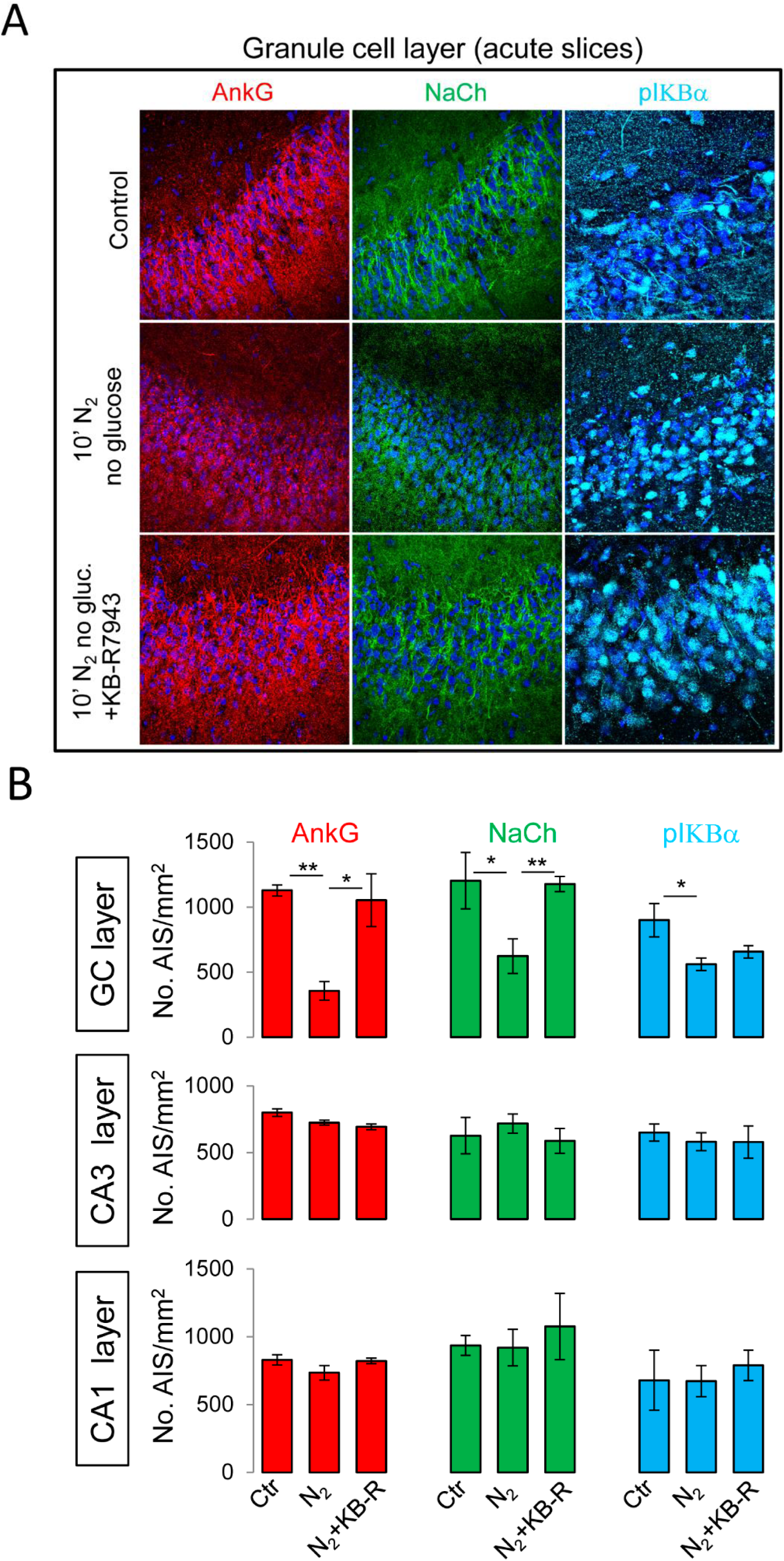
AISs are quickly dismantled specifically from granule cells after brief oxygen deprivation in acute slices. Blockade of NCX_rev_ with KB-R7943 prevents AIS dismantling. (**A**) GC layer immunostaining for ankyrin-G, sodium channel (panNaCh) and pIKBα in acute hippocampal slices in three different conditions: (i) control, (ii) oxygen and glucose deprivation (OGD; 10 min N_2_, no glucose) and (iii) OGD+KB-R7943 (2μM), as indicated in the figure. (**B**) Average number of AISs for each experimental condition in GCs, CA3 and CA1 layers (*images not shown for CA1 and CA3 layers)*. Data are mean ± SEM from 4 animals for each immunostaining. Four slices per animal were averaged. *p<0.05,**p<0.01

AIS disassembly should increase cell firing threshold, thus reducing cell excitability (Colbert & Johnston, 1996; Zhou *et al*., 1998; Kole *et al*., 2008; Winkels *et al*., 2009). We measured AP threshold after OGD in GCs from slices treated with KB-R7943 or vehicle (both washed-out for >20 min prior recording to exclude acute effects of the drug). The firing threshold was a 17±5% (n = 9, p < 0.05) higher in control conditions −disrupted AISs−, than after KB-R7943 treatment −protected AISs− (Fig.4A-B). Moreover, the number of cell-attached-recorded spikes, evoked by perforant-path stimulation, was gradually reduced in control slices over increasing time post-OGD whereas KB-R7943 treatment largely abolished this decay (Fig.4D). In both conditions synaptic inputs onto GC layer (measured as fEPSPs amplitudes nearby the patched GCs) where similarly preserved (Fig. 4E). These results suggest that NCX-mediated disassembly of AISs after OGD increases AP threshold thus reducing GC excitability.

**Figure 4.**
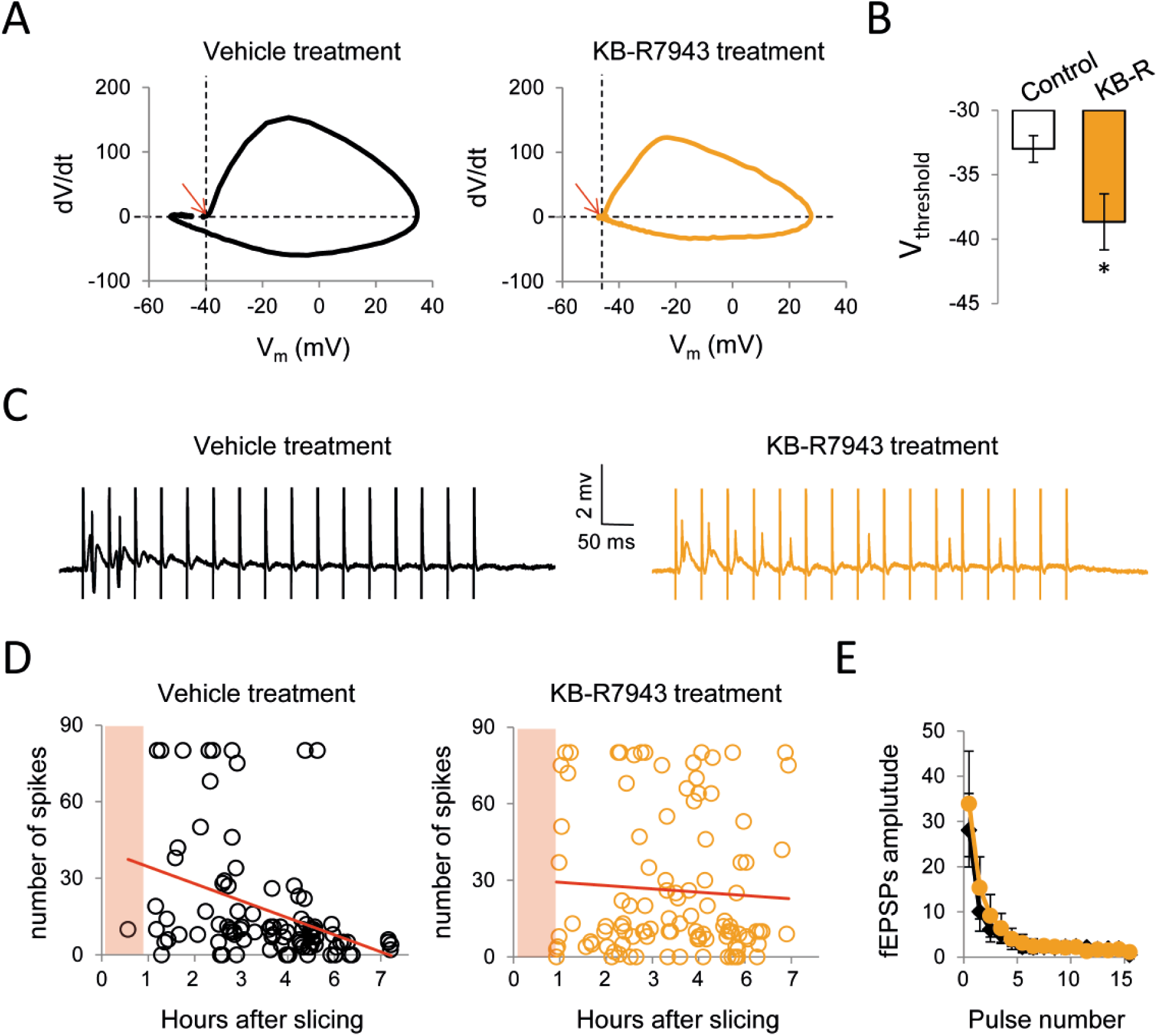
Prevention of AIS dismantling by NCX_rev_ blockade preserves GC excitability during O_2_ deprivation in acute slices. (**A**) Phase plots of APs recorded in whole-cell from GCs in slices exposed to vehicle (black) or KB-R7943 (orange) during OGD for 1h. Recordings started after wash-out of the drug for >20 min. Red arrows indicate firing threshold. (**B**) Effect of KB-R7943 (2μM) on firing threshold (n=9 for each condition). (**C**) Cell-attached recordings of APs from GCs synaptically activated by perforant path stimulation with 5 trains of 16 pulses at 20 Hz (one train every 10 seconds). (**D**) Number of spikes elicited by the mentioned stimulating protocol for all recorded cells at different times after slicing. Each dot represents one cell. The pink shadow indicates the window of OGD. Black dots correspond to vehicle treated slices. Orange dots correspond to KB-R7943 treated slices. Red lines are fitted lines to the data. Slopes are significantly different; p<0.015. (**E**) Amplitude of field EPSPs recorded at the GC layer in the presence in KB-R7943/vehicle treated slices. Values are mean ± S.E.M, ** p<0.01; *p<0.05

### AIS disruption after brief ischemia *in vivo*

*In vivo,* we observed a similar AIS disruption than upon OGD in acute slices. A brief global transient ischemia (see Methods) produced a dramatic disappearance of ankyrin-G (and, more moderately, of pIκΒα) in the GC layer, after just 30 min reperfusion (Fig.5A-B). Instead, Na^+^ channels where not affected after 30 min, but they vanished after 1 day of recovery from ischemia.

**Figure 5.**
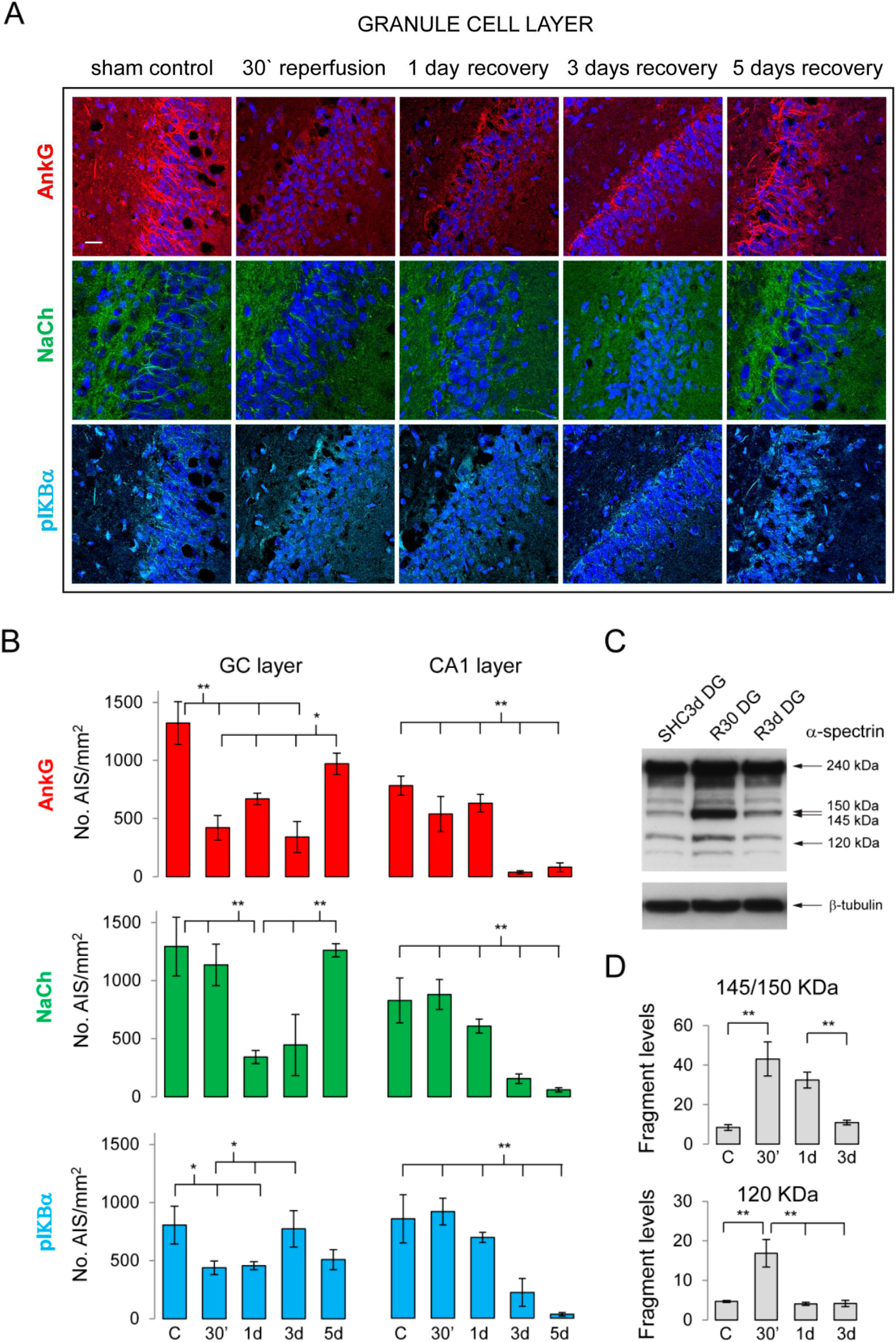
Brief ischemia *in vivo* induces reversible AIS dismantling and reversible calpain activation in the dentate gyrus, but not in CA1 area. (**A**) Immunostaining for Ankyrin G, panNaCh and pIκΒα in control pre-operated animals (sham) and in rats following variable periods of reperfusion (30 min, 1 day, 3 days and 5 days) after ischemia. (**B**) Summary of the results for the number of AIS in GCs and CA1 area for all periods of recovery. The pictures for CA1 layer are not shown (see Supplementary Fig.5). Values are averages of 4 rats for each condition ± SEM. Scale bar=30μm. (**C**) Western blot showing α-spectrin degradation by dentate gyrus calpain/caspase-3 enzymatic activities in control (sham), 30 min (R30) and 3 days (R3d) of reperfusion after ischemia. The molecular weight of the digested fragments is indicated. (**D**) Average of 145/150kDa (calpain related) and 120kDa (caspase-3 related) spectrin fragments from 4 experiments of each condition (control: C; 30 min: 30’; 3 days: 3d). *p<0.05; **p<0.01

To our surprise, after 5 days of recovery, AISs immunostaining in GC layer was almost fully recovered (Fig.5A-B). In contrast to the effect of ischemia on GCs, in CA1 area the three AIS markers remained intact for at least 1 day after ischemia. At day 3, they finally disappeared from CA1, without recovery (Fig. 5B and Supplemental Fig. 5). CA3 area suffered an intermediate situation: a mild reduction in the number of AISs at day 1, that was not recovered (Supplemental Fig. 6).

As calpain is responsible for irreversible AIS disruption (Schafer *et al*., 2009), we tested if *reversible* AIS dismantling in GCs was also induced by calpain (Fig. 5C-D). We studied calpain activity in isolated dentate gyrus (DG) by measuring the specific spectrin breakdown product of 145/150-kDa, as well as the 120-kDa caspase-3 mediated spectrin fragment, a marker of apoptosis (Nath *et al*., 1996). Calpain and caspase-3 activities were increased after 30min of reperfusion from ischemia (Fig. 5C-D), coinciding with Ankyrin-G dismantling in GCs (Fig. 5B). Moreover, after 5 days of reperfusion −when AISs are fully reconstructed in control conditions− calpain activity returns to normal values. This suggests that calpain mediates GC-AIS disruption, and then reverses its activity to basal conditions, allowing reconstruction of the AIS.

In CA1 area, where AISs remain intact for at least 48-fold longer, calpain activity was moderately elevated, but increased dramatically after 3 days from ischemia, coinciding with irreversible AIS dismantling in these cells. In contrast to GCs, CA1 caspase-3 was also elevated at day 3, indicating a path towards apoptosis in this region (Supplemental Fig. 5C-D).

We next studied the role of NCX_rev_ on AIS dismantling *in vivo*, by comparing the number of AISs in the GC layer in rats injected with KB-R7943 or vehicle prior ischemia induction. After 30 minutes of reperfusion, ankyrin-G was highly preserved in KB-R7943 treated animals, as compared to vehicle treatment (Fig. 6A-B). CA3 and CA1 areas displayed no changes in their amount of AISs with KB-R7943 (Supplemental Fig. 7). After 1 day of recovery, the protection by the drug in GC layer was extended to the three AIS markers (Fig. 6B). After 5 days, GCs showed no changes with the drug, except for pIκΒα antibody, that labelled a higher number of AISs (Fig. 6B). Immunolabeling of pIκΒα reflects phosphorylation levels rather than activation of the NFκΒ pathway (Herkenham *et al*., 2011; Buffington *et al*., 2012), thus low levels of AIS-pIκΒα could indicate energy deficiency and/or represent a signal for AIS *assembly*, a ‘call’ for reconstruction (Konig *et al*., 2017).

**Figure 6.**
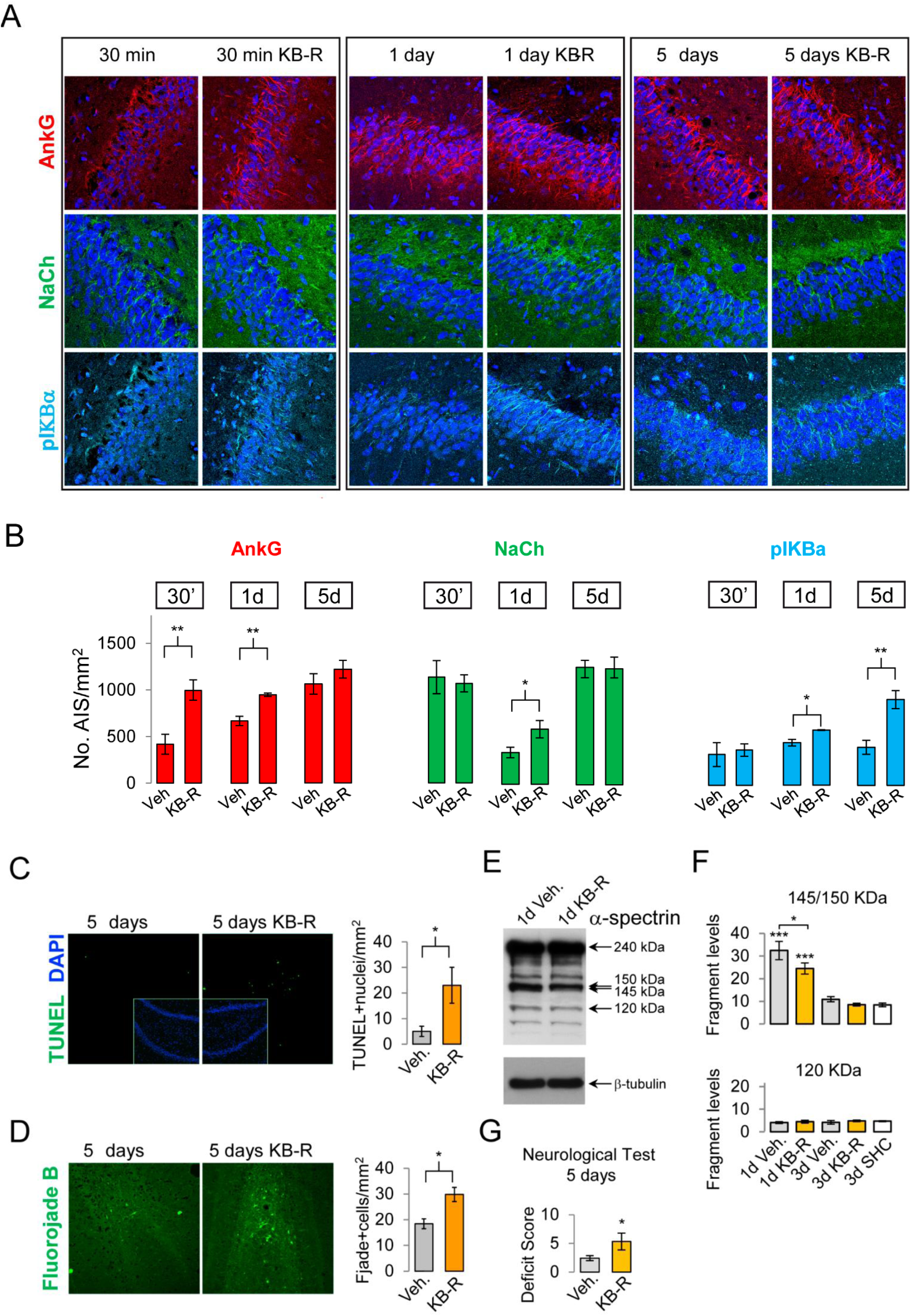
NCX blockade *in vivo* preserves AISs in GCs but is harmful for neurons in the hilus. (**A**) Effect of KB-R7943 or vehicle injection before ischemia on AIS integrity (Ankyrin-G, panNaCh and pIKBα) at 30 min, 1 day and 5 days reperfusion following ischemia. (**B**) Summary of results for the number of AISs in vehicle and KB-R7943 treated rats. Data are averages ± SEM of n=4 rats per condition. (**C**) TUNEL staining of the dentate cells (the inset shows dapi staining) 5 days after ischemia in rats treated with KB-R7943 or vehicle. *Right*: summary of the results for 4 animals per condition. (**D**) Fluorojade B staining in the dentate gyrus from the same tissue as (C). *Right*: summary of the results. Data are mean ± SEM of n=4 rats per condition. (**E**) Blot showing α-spectrin degradation by dentate gyrus calpain/caspase-3 activities from rats injected with KB-R7943 or vehicle (veh.) before ischemia and reperfusion for 1 day (R1d). The apparent molecular weight of spectrin and of the breakdown products is indicated. (**F**) Summary of the data for 145/150-kDa and 120-kDa from 4 rats per condition. *p<0.05; **p<0.01; ***p<0.001; n=4 rats per condition (**G**) Effect of KB-R7943 on neurodeficit score outcomes. The bar graphs show the neurological Deficit Score in animals treated with KB-R7943 (KB-R) or vehicle (Veh) at 5 days of reperfusion after ischemia. *p < 0.05, from 7 to 10 rats per condition.

To determine if AIS protection by NCX blockade modifies cell death rate after ischemia, we quantified apoptosis with TUNEL assay in KB-R7943 or vehicle treated rats, after 5 days from ischemia (Fig.6C). GCs are highly robust against ischemia, as opposed to CA1 area. We did not observe cell death in the GC layer either in control or in KB-R7943 treated animals. However, KB-R7943 treated animals showed a 460±140% increase in cell death rate in the hilus (n=4; p<0.05), as compared to control (Fig. 6C). Intriguingly, it is reported that overexcitation of GCs leads to damage of their targets in the hilus (Hsu & Buzsaki, 1993; Jolkkonen *et al*., 1997; Norwood *et al*., 2011). In CA1 area, whose AISs were partially protected by KB-R7943 (at day 5), the drug increased cell death (20%±9%, n=4, p<0.05; Supplemental Fig. 8).

We have also used a more general marker of cell damage: Fluorojade-B staining (Methods). The number of Fluorojade-B stained cells in the hilus was higher in KB-R7943 treated animals (day 5, n=4, p<0.05; Fig.6D), consistent with TUNEL data. Ischemia also increased Fluorojade-B stained cells in CA3 area. However, KB-R7943 had no significant effect on that (8.88±1.99, control; 11.61±4.36, drug; cells/mm^2^; n=4). And we also tested Fluorojade-B staining in acute slices. In contrast to *in vivo* staining, OGD deprivation highly increased Fluorojade staining of the GC layer, and KB-R7943 worsened this damage (n=4, p<0.05; Supplemental Fig. 9). If, indeed, NCX mediated Ca^2+^-entry activates calpain *in vivo*, then KB-R7943 should decrease calpain activity after ischemia. We therefore studied the effect of KB-R7943 on calpain/caspase-3 activities in the DG. KB-R7943 reduced by 25±7% the DG calpain-dependent 145/150-kDa-spectrin fragment after 1 day from ischemia (n=4; p<0.05; Fig.6E-F). Interestingly, despite the parallelism of calpain and caspase-3 activities after ischemia in the DG (Fig.5D), only calpain activity but not caspase-3 was reduced in KB-R7943 treated animals (Fig 6E-F), indicating that Ca^2+^ entry through NCX_rev_ specifically activates calpain. Importantly, consistent with our cell survival assays, KB-R7943 treated animals performed significantly worse the neurological tests (Fig.6G). In contrast, in CA1 area KB-R7943 did not affect calpain or caspase-3 activities for at least 1 day after ischemia (Suplementary Fig.7G). At day 3, CA1-calpain activity but not CA1-caspase-3 activity was reduced by the drug (Supplementary Fig. 7F-G) which could explain the preservation of AISs observed at day 5 in CA1 area (Supplementary Fig.7D). In fact, ischemia produces very high frequency firing in CA1 area (Barth & Mody, 2011), which could explain why drug protection of AISs increases cell death in this region (Supplementary Fig. 8). We did not observe apoptotic death in CA3 area after ischemia (not shown), although delayed death has been reported in this area after longer ischemia (Wang *et al*., 2004).

**Figure 7.**
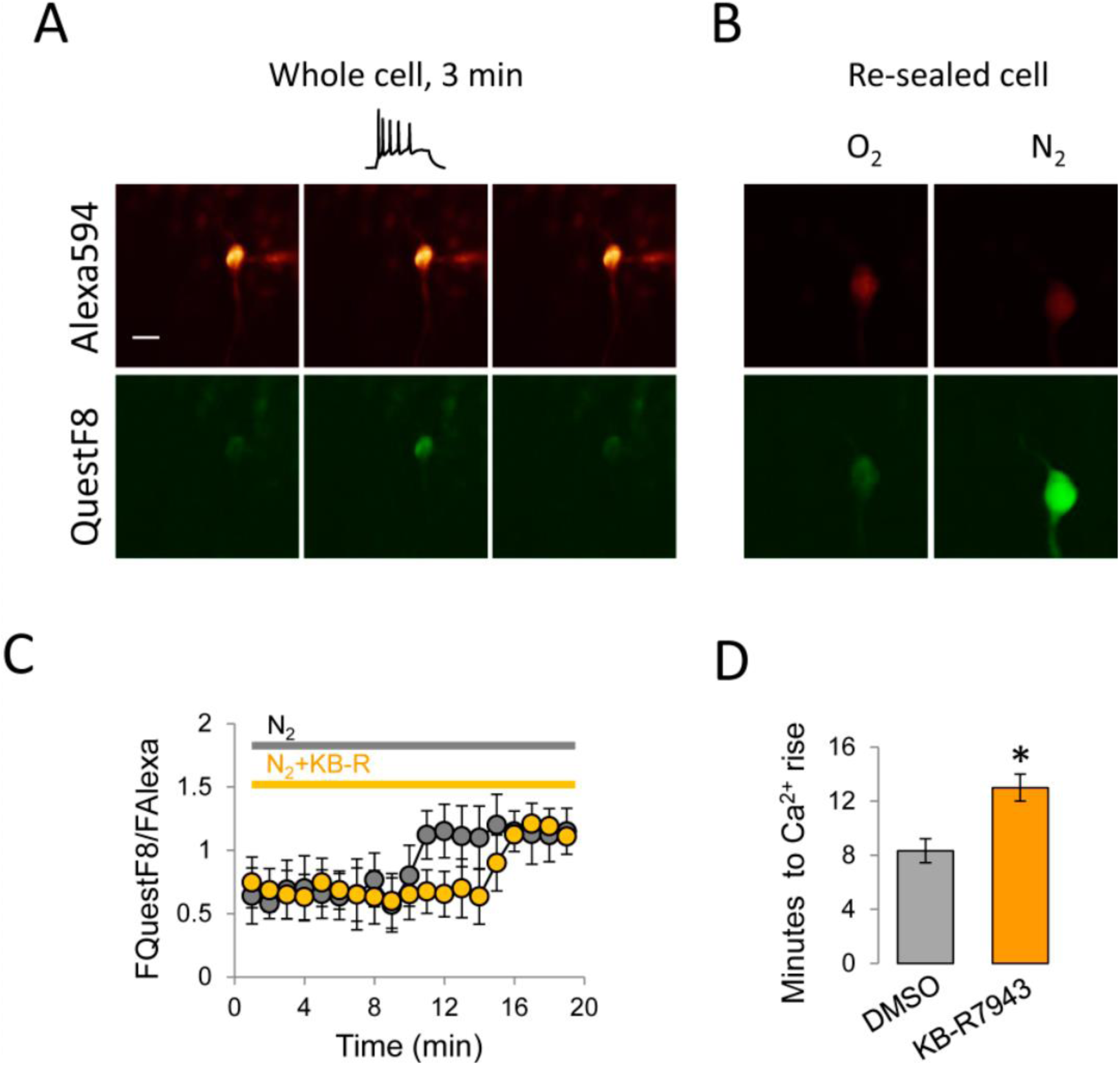
Na^+^/Ca^2+^ exchanger blockade delays granule cell Ca^2+^ rise induced by OGD in acute slices. (**A**) Example of a granule cell patched in whole-cell configuration after 3 minutes from breaking in. The pipette contained 400μM Quest-F8 and 80μM alexa594. (**B**) Alexa594 and Quest-F8 fluorescence before and after OGD at 32°C. A global increase of internal calcium is observed in the green channel. Image recording was every 30 seconds during the first 6 minutes after starting N_2_ ringer perfusion, and every 10 seconds until the end of the experiment. (**C**) Time course of QuestFluo8Ca^2+^/Alexa594 signal in cells exposed to the vehicle (*grey*) or KB-R7943 (2μM, *orange*). (**D**) Average time to Ca^2+^ rise from the start of N_2_ Ringer with/without the drug. Data are mean ± SEM of 9 cells per condition; p<0.025. Scale bar is 15 μm.

The mechanism of AIS disruption for longer ischemia depends on an increase of intracellular calcium (Schafer *et al*., 2009), that would then activate calpain (Moldoveanu *et al*., 2002). To investigate if NCX_rev_ introduces Ca^2+^ into granule cells during OGD, we monitored resting Ca^2+^ in cells loaded with Quest-Fluo8 and Alexa Fluor-594 (Fig.7). In control conditions, resting Ca^2+^ increased after 7-8 minutes of OGD. Interestingly, with KB-R7943 in the bath, Ca^2+^ rise was delayed by 53±12% (Fig.7C&D), thus suggesting a Ca^2+^ entry through NCX_rev_ during ischemic stress.

## DISCUSSION

We have found that NCX operating in the reverse mode (NCX_rev_, Na^+^ efflux mode) contributes significantly to the Na^+^ extrusion and Ca^2+^ influx at the AIS during short, physiologically relevant, trains of spikes. This mechanism seems to act exclusively at the AIS. The reverse mode of the NCX has also been reported to occur in synapses, although its recruitment requires stronger repetitive of high frequency stimulation (Minami *et al*., 2007; Roome *et al*., 2013)

Our experimental observations implicating NCX_rev_ during repetitive spiking appears consistent with the notion that (i) the reverse NCX mode is favoured by depolarization (Blaustein & Lederer, 1999); (ii) Na^+^ channels activate especially fast at the AIS (Schmidt-Hieber & Bischofberger, 2010) during depolarization while the spike dependent Ca^2+^ entry mainly occurs during the repolarising phase of the AP (Borst & Sakmann, 1998); (iii) free-Ca^2+^ is quickly bound by endogenous buffers while Na^+^ is not; (iv) the low micromolar range of Ca^2+^ is required for NCX reverse mode activation (Blaustein & Lederer, 1999); and (v) average Na^+^ influx at the AIS is the largest across the neuron, 0.4-1mM/spike (Shu *et al*., 2006; Scott *et al*., 2014). In addition, the possibility that the NCX could closely colocalize with Na^+^ channels at the AIS has to be taken into account. Notably, the NCX reverse mode has been reported to occur in presynaptic terminals upon strong stimulation (Regehr, 1997; Roome *et al*., 2013), and this mode of operation has been experimentally shown and modelled under relatively mild stimulation protocols (Minami *et al*., 2007). Again, compared to presynaptic terminals, the AIS displays much greater Na^+^ influx and lower Ca^2+^ influx during spikes trains.

Despite the potent effect of NCX on Na^+^/Ca^2+^ interplay at the AIS during short burst of activity, its effect on spike generation is relatively mild in conditions of healthy concentration of internal Na^+^. This led us to the hypothesis that the AIS-NCX might be important in the longer term, and especially under metabolic stress, a situation that raises the levels of internal Na^+^ due to the progressive loss of membrane potential control, which activates low threshold/persistent current through Na^+^ channels (Stys *et al*., 1992; Stys & Steffensen, 1996; Craner *et al*., 2004). In fact, we show here that subthreshold constant depolarizations produce AIS Na^+^ loads equivalent to long trains of high frequency action potentials in granule cells.

The role of NCX in neurodegeneration has been under debate. It seems clear that NCX blockade protects myelinated axons from degenerating conditions (Petrescu *et al*., 2007; Norwood *et al*., 2011; Barsukova *et al*., 2012; Bei & Smith, 2012). Intriguingly, while NCX2 and NCX3 knockout mice are more sensitive to damage by long duration (90-120 min) focal ischemia (Jeon *et al*., 2008; Molinaro *et al*., 2008), acute blockade of NCX with the specific blocker SEA0400 ameliorates the impact of similar ischemic conditions (Wang *et al*., 2007). Our results are consistent with a protective action of NCX reverse mode on axonal properties. However, the overall long-term effect is more complicated. Based on our data, we propose the following model (Fig 8): during brief ischemia the exchanger contributes to a calcium signal, likely originating at the AIS. Ca^2+^ rise would promote calpain/caspase-3 activity. These enzymes would quickly dismantle AISs, raising the firing threshold. Lowered cell excitability should thus prevent dangerous overexcitation of the trisynaptic hippocampal circuit during harmful stress. For longer ischemia, AIS dismantling would not be protective and would lead to neural degeneration (Schafer *et al*., 2009). Instead, for short ischemia, after oxygen reperfusion, the replenishment of energy would allow the cell to return to normal Ca^2+^ levels (by engaging Ca^2+^ pumps) allowing the calpain activity to return to basal values. In these conditions, AISs would start being reconstructed and a few days later, granule cells would be recovered. An intriguing possibility is that without the AIS being dismantled inside a brief critical period, hippocampal damage would be higher due to overexcitation of the GC-CA3 and GC-hilus synaptic pathways.

**Figure 8.**
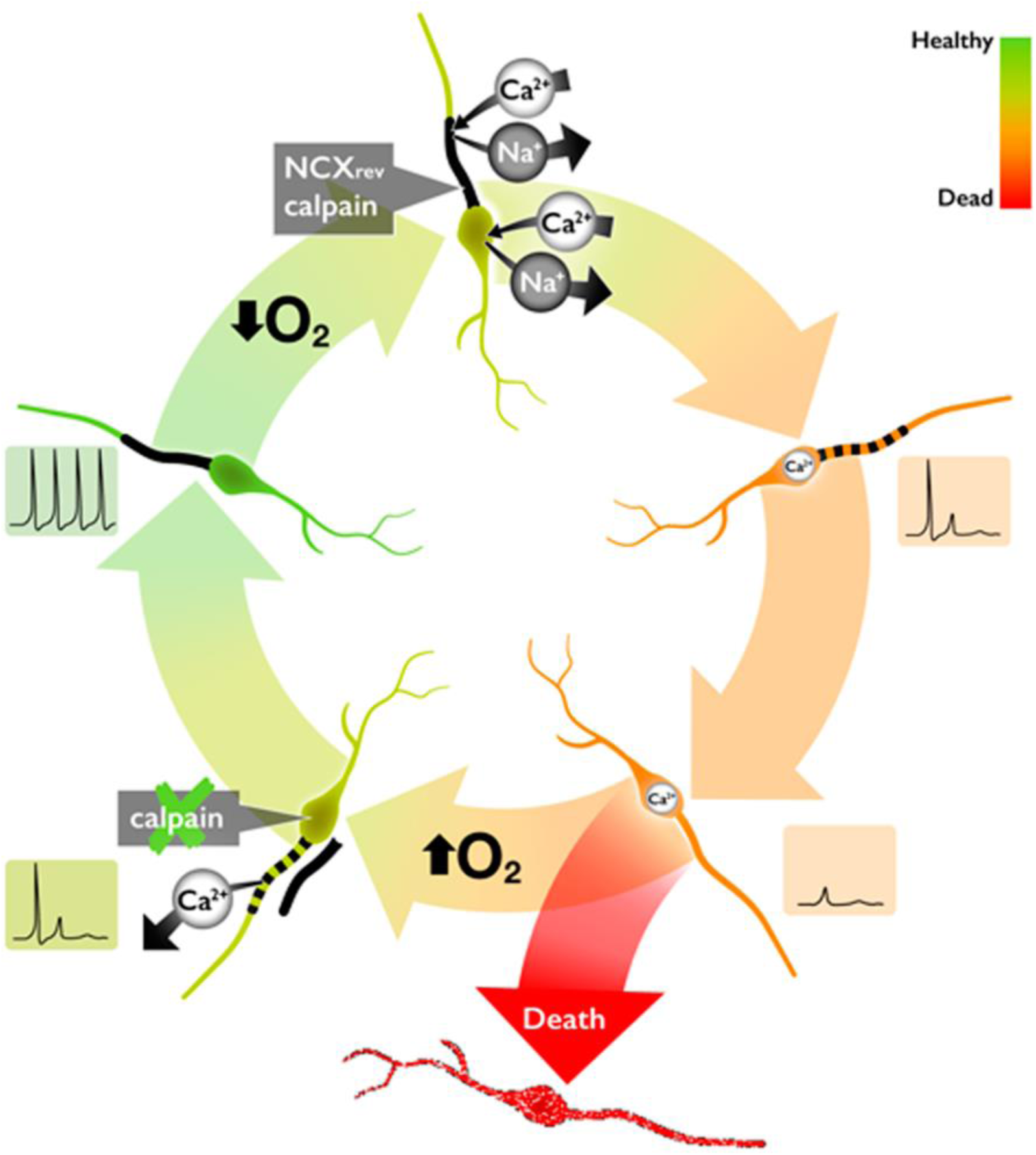
Proposed dynamics of AIS disassembly after brief ischemia. The scheme starting point is a healthy granule cell (*left* part of the scheme, in *green*) able to fire APs normally. The AIS is intact and functional (*black thick line*). Then, ischemia starts and O_2_ levels are reduced. Energy exhaustion leads to a mild depolarization that produces Na^+^ entry, most prominently in the AIS (as observed; not show in the scheme). The NCX removes the excess of Na^+^ introducing Ca^2+^ into the cell. This Ca^2+^ activates calpain, responsible of proteolysis of ankyrin-G from the AIS (dotted thick line), so that sodium channels start detaching from the AIS and diffusing along the membrane (as observed; not shown in the scheme). The AIS disassembly reduces granule cell excitability (as observed) avoiding hippocampal damage, to the hilus (as observed). If oxygen is re-established soon enough (short ischemia), the silent cell can extrude the excess of Ca^2+^ so that calpain activity would revert (as observed), and AIS could be reconstructed (as observed). The cell would then return to normality without damage, and so its targets (hilus). If AIS dismantling did *not* occur quickly (as observed with NCX blockade) the damage would be greater (as observed), presumably due to overexcitation. This mechanism could also contribute to the very low mortality of granule cells and their connected cells after brief ischemia.

A reversible disassembly can be of course interpreted simply as a not sufficient stress for the cell, which can recover after some time, rather than a designed mechanism to improve survival. However, the reversibility of calpain/caspase-3 activities has not been reported before despite the numerous studies dedicated to the topic. This suggests that a specific mechanism of reversion could be involved in the reversible AIS dismantling reported here. In fact, recently, AISs and nodes of Ranvier have been reported to recover from crushed optic nerves (Marin & Rasband, 2017).

The observed hilar damage after transient ischemia in KB-R7943 treated rats could be explained by a direct effect of KB-R7943 in hilar cells rather than by GC firing. However, KB-R7943 preferentially blocks NCX3 transporters (Iwamoto & Shigekawa, 1998), and hilar cells do not express NCX3, while GCs express them abundantly (Canitano *et al*., 2002). During brief ischemia GCs are barely damaged. Thus, without a brake these cells would persistently fire and discharge for a long time, damaging their targets.

The hilus is densely contacted by GC mossy fibers. Importantly, GC overactivity is reported to damage the hilus (Jolkkonen *et al*., 1997; Norwood *et al*., 2011). In fact, mossy fiber synaptic targets, including the hilus, are highly vulnerable during ischemia (Hsu & Buzsaki, 1993). Therefore, the damage we observe in the hilus in KB-R7943 treated animals could be due to a higher excitability of GCs after the ischemia. In fact, GC firing threshold is reported to be reduced after ischemia *in vivo*, while spontaneous activity is not altered (in the absence of NCX blockers) (Gao *et al*., 1998).

Finally, the timing of appearance of new mature neurons or axonal sprouting after ischemia in the DG (Liu *et al*., 1998; Kee *et al*., 2001; Yagita *et al*., 2001; Kawai *et al*., 2004; Hinman *et al*., 2013) is not consistent with the timing of recovery of AISs we observed after 5 days from global transient ischemia.

This extensive work reveals a novel pathway for Ca^2+^ to enter the AIS, the NCX, that contributes to preserve normal firing properties of GCs. In addition, we find that NCX triggers AIS disassembly during brief ischemia by activating the protease calpain; and that this process is fully reversibly after a brief period of recovery.

## EXPERIMENTAL PROCEDURES

### Electrophysiology

Acute 350 μm hippocampal slices from three- to four-week old male Sprague-Dawley rats were transferred to a submersion-type recording chamber (Scientific Systems Design, NJ), and superfused with (mM): 124 NaCl, 2.5 KCl, 2 CaCl_2_, 1 MgCl_2_, 26 NaHCO_3_, 10 glucose and bubbled with 95% O_2_ / 5% CO_2_. The slice orientation that allows intact mossy fibers (MFs) to be preserved was detailed previously (Scott & Rusakov, 2006). The internal solution included (mM): 135 K methanesulfonate, 2 MgCl_2_ 10 HEPES, 10 Na_2_-Phosphocreatine or 10 Tris-phosphocreatine (according to final [Na^+^]_i_), 4 Na_2_ATP, 0.4 NaGTP, and fluorophores as indicated. Unless specified otherwise, dentate granule cells were held at −80 mV; cells were discarded if their resting potential, in current-clamp mode, increased above −70 mV during the experiment. Single orthodromic escape action currents were evoked by a 2 ms command voltage pulse. Recording sweeps (normally 500 ms long) were collected at 5 kHz with 30 s or 1 min intervals. Experiments were carried at 22-24°C unless specified otherwise. Receptor antagonists were purchased from Tocris Cookson, and Alexa-594, Fluo-4 and SBFI were purchased from Invitrogen; QuestFluo-8 from Teflabs (US). All animal handling procedures followed current EU regulations.

### Two-photon excitation fluorescence imaging

We used a confocal scanning unit (LCS Leica SP5 II, Leica Microsystems, Mannheim, Germany) attached to an upright microscope (DM6000CFS, Leica; 25x objective, HCX IRAPO L 25x, numerical aperture 0.95, Leica) linked to a MaiTai laser (in Alicante), or a Radiance 2000 or 2100 (Zeiss-BioRad) imaging system optically linked to a MaiTai (SpectraPhysics-Newport) femtosecond pulsed infrared laser (in London). Granule cells held in whole-cell mode were loaded with three types of fluorophores: a morphological tracer Alexa Fluor 594 (20-40 µM); the high-affinity Ca^2+^ indicators, Fluo-4 or Quest-Fluo8 (200 μM); and the low affinity Na^+^ indicator SBFI (1mM) as indicated. Fluorophores were excited in two-photon mode at 800 nm or 760 nm respectively. The axon was imaged at the maximal optical resolution (∼0.2 μm, 70 nm per pixel).

The Ca^2+^ dependent Fluo-4 or QuestFluo-8, and the Na^+^ dependent SBFI fluorescence signals were quantified as ΔF/F_B_ = (F – F_pre_) / (F_pre_ – F_0_), where F_0_ denotes the background fluorescence and F_B_ = F_pre_ – F_0_ the baseline axonal fluorescence before any stimulus was delivered (or Ca^2+^ rise occurred after OGD). For Ca^2+^ transients evoked by precisely-timed spikes generated in response to brief depolarizing pulses (Fig. 1B *right*), the ratio of Ca^2+^ - dependent fluorescence increments during the spike train is given as ΔF_3_/ΔF_1_ for the third and first spikes. When Ca^2+^ transients were elicited by depolarization-induced spike trains (Fig. 1B *right*), the ratio of early fluorescence increments F_early_ (0-40 ms after stimulus onset) and delayed increments F_late_ (40-200 ms after stimulus onset) was calculated: (F_late_ – F_B_) / (F_early_ – F_B_).

Image analysis was performed on stacks of stored line-scan images using *ImageJ* macros. False color tables and averaged images were used for illustration purposes but the quantitative analyses always dealt with the original (gray level) pixel brightness values. In most experiments, we reconstructed the axon trajectory using a collage of high-resolution Kalman-filtered z-stacks 15-20 µm deep using 10-scan average frames.

### Compartmental NEURON model of a dentate granule cell

As previously described (Scott *et al*., 2014), we used a NEURON model of a fully reconstructed dentate granule cell adapted from Schmidt-Hieber (2007), cell No 7, imported from SenseLab, ModelDB=95960 at http://senselab.med.yale.edu/modeldb (Schmidt-Hieber *et al*., 2007). Briefly, passive axon parameters were adopted from measurements made earlier (Scott *et al*., 2008). Axonal axial resistance was set to 80 Ohm·cm. Sodium and channel models where implemented as described(Schmidt-Hieber *et al*., 2008) and distributed as a function of distance from the soma. The channel distribution was optimized to reproduce experimental observations (without introducing additional conductances). The reverse potentials for Na^+^, K^+^, and leak currents were set to 58, -95 and -80 mV, respectively. Na^+^ concentrations changes were simulated by manipulating reversal potential for Na^+^. Action potentials were elicited by somatic current injections as illustrated.

### Oxygen and glucose deprivation (OGD) in acute slices

Acute slices were perfused for 10 minutes with a Ringer solution nitrogenated with a mixture of 95% Nitrogen + 5%CO2 without glucose at the same osmolarity as normal Ringer (300mOs). For some experiments, OGD was simply applied by stopping oxygenation (maintaining pH7.3 with 5mM EGTA), perfusing Ringer without glucose, and rising [K^+^] to 14mM for 1h. After these periods of OGD, slices were transferred on an interface chamber with normal oxygenated Ringer for recovery for at least 20 minutes.

### Immunohistochemistry

After treatments, acute slices were fixed in 4% PFA for 2 hours and then washed out with PBS. For immunodetection, slices were treated with 50 mM NH4Cl and incubated in blocking buffer for 45 min (PBS, 0.22% gelatin, 0.1% Triton X-100). After blocking non-specific binding, slices were incubated for 1 h at room temperature with the primary antibodies diluted in blocking buffer. The primary antibodies used were: rabbit anti-phospho IκBα (1:1000, Cell Signaling), mouse anti-ankyrinG (1:500, Sigma); mouse anti-PanNaCh (1:500, Sigma-Aldrich). The secondary antibodies were donkey anti-mouse, anti-rabbit or antigoat coupled to Alexa-488, Alexa-594 or Alexa-647 (Invitrogen). After staining, the coverslips were mounted in Fluoromount G (Southern Biotech.)

For *in vivo* experiments, animals were killed by transcardial perfusion performed under deep anesthesia. Perfusion via the left ventricle was started with a washout of 200 ml of 0.9% NaCl and the brains, following perfusion and fixing with 4% (w/v) paraformaldehyde solution in PBS, were removed and postfixed in the same solution overnight at 4 °C. Brains were washed sequentially with 10, 20 and 30% (w/v) sucrose in PBS, embedded in Tissue-Tek O.C.T. (Sakura Finetek) and frozen at -80 °C prior to cryostat sectioning. Brain coronal sections (5-20 μm thick) were prepared at the level of interaural +5.7 ± 0.2 mm on Real Capillary Gap microscope slides (Dako).

### AIS quantification

Images were obtained using a laser scanning spectral confocal microscope Olympus Fluoview FV1200 (Japan). The number of AIS from at least 10 fields from 3-4 slices per condition was counted double-blinded in Fluoview software (Olympus, Japan). AIS count in the GC layer labelled with panNaCh-Ab, AnkG and pIκBα was performed with z-staks of confocal images To identify false positives, the sequence of a z-stack of the same area was visualized in a continuous mode (as a movie). Each of these images was counted blindly and in random order by 2 different persons. The final result was the average of both measurements. All images shown and analyzed throughout the paper correspond to single confocal planes obtained at the same depth and with the same optical parameters for all slices (Supplementary Fig.10).

We also used Image J (NIH, US) automatized analysis of the total area labelled with Ankyrin-G, obtaining similar results. However, we preferred the visual double-blind analysis because AISs in GC layer display frequent crossover. In addition, panNaCh-Ab, and specially pIκΒα-Ab, also produce somatic staining. These issues complicated automated counting of AISs in the GC layer.

### Animal model of ischemia

Transient forebrain ischemia was induced in adult Wistar rats (10-12 weeks, from Charles River) by the standard four-vessel occlusion model described previously (Garcia-Bonilla *et al*., 2007; Ayuso *et al*., 2010; Ayuso *et al*., 2013). Briefly, both vertebral arteries were irreversibly occluded by electrocoagulation under anesthesia with a mixture of atropine, ketamine and diazepam (0.25, 62.5, and 5 mg/kg, respectively) delivered by intra-peritoneal injection. After 24 h, ischemia was induced by carotid occlusion with small atraumatic clips for 15 min and then clips were removed from the carotid arteries for reperfusion. After 30 min, 1, 3 or 5 days of reperfusion (R30, R1d, R3d and R5d, respectively), the animals were sacrificed. Sham control (SHC, SHC3d or SHC5d) animals were prepared in the same way as the R30, R3d or R5d animals, respectively, but without carotid occlusion. In some experiments, animals were treated with 8 mg/kg KB-R, diluted in 1.2%-12% DMSO-ethanol in saline (v/v) as vehicle, by intra-peritoneal injection 45 min before ischemia induction. The treatments were blindly performed, assigning a random order to each vehicle or treated animal. All procedures associated with animal experiments were approved by The Ethics Committee of the Hospital Ramon y Cajal, Madrid, Spain, and in accordance with the ARRIVE (Animal Research: Reporting In Vivo Experiments) guidelines.

### Evaluation of neurological deficits

The neurological deficits in rats subjected to ischemia were evaluated blindly using a validated scale described in Geocadin et al.(2000). The neurological deficit score was determined in R5d animals to evaluate the effects of KB-R7943 following ischemia-reperfusion. The score ranges from 0 (best) to 10 (rats did not walk spontaneously and had a depressed level of consciousness) and it includes a score of general behavioral deficit, subscore in motor assessment, and sensory assessment subscore.

### Western blot analysis

Mouse monoclonal anti-α-spectrin antibody was from Chemicon. Dentate gyrus from control and ischemic animals were rapidly dissected. The samples were homogenized and processed to obtain a postmitochondrial supernatant (PMS) as described in ref (Ayuso *et al*., 2013). All procedures were performed at 4 °C. The sample corresponding to each animal was separately kept at -80 °C until used and protein concentrations were determined for each sample.

Samples of PMS (35 μg) of each different experimental condition were analyzed by SDS-PAGE (7.5% acrylamide, 2.6 % crosslinking), transferred onto PVDF membranes (GE Healthcare), incubated with the primary antibody against the specific protein, and after the blots were incubated with peroxidase-conjugated anti-mouse IgG (GE Healthcare), and developed with ECL reagent (GE Healthcare). The western blots were quantified using Quantity-One software (Bio-Rad). Internal standards (tubulin) were included to normalize the different immunoblots.

### Statistical analysis

The different animals from each experimental condition or group were independently analyzed in duplicate and their averaged values were used for statistical analysis. Data were from 3 to 7 different animals run in duplicate. Statistical analysis was performed using analysis of variance (anova) following Newman-Keuls’ post test, when analysis of variance was significant, to compare the data between experimental groups. Comparisons between two groups were done by Student’s *t*-test. Statistical significance was set at p < 0.05 using Prism statistical software (GraphPad Software).

## Acknowledgements

Ignacio Lizasoaín, Juan Lerma, Félix Viana, Marco Capogna and Rubén Deogracias for carefully reading the manuscript. Hanoch Kaphzan and Christine Ross for useful discussion. Stuart Bailey for illustration assistance.

This work was supported by a Ministerio de Ciencia e Innovación Grant to RS (SAF2010-200604), NRW-Rückkehrerprogramm, Human Frontiers

Science Program, German Research Foundation (SFB1089, SPP1757), DAAD, UCL Excellence Program to CH.

## Author Contributions

CS performed all immunohistochemistry, cell survival assays, in vitro degeneration experiments and analysis.

EMA, MIA performed ischemia, spectrin assays and TUNEL. AA ischemia experiments. AA and RS analyzed ischemia experiments data.

RS performed most electrophysiology and imaging experiments and analyzed the data. CH performed some modelling, imaging experiments and analysis. PR performed some imaging and electrophysiology experiments, and analysis.

RS conceived the project, designed experiments, and wrote the manuscript to which all authors contributed.

