## Supplemental Material for "Na^+^/Ca^2+^ exchanger triggers transient disruption of axon initial segments in hippocampal granule cells after brief ischemia"

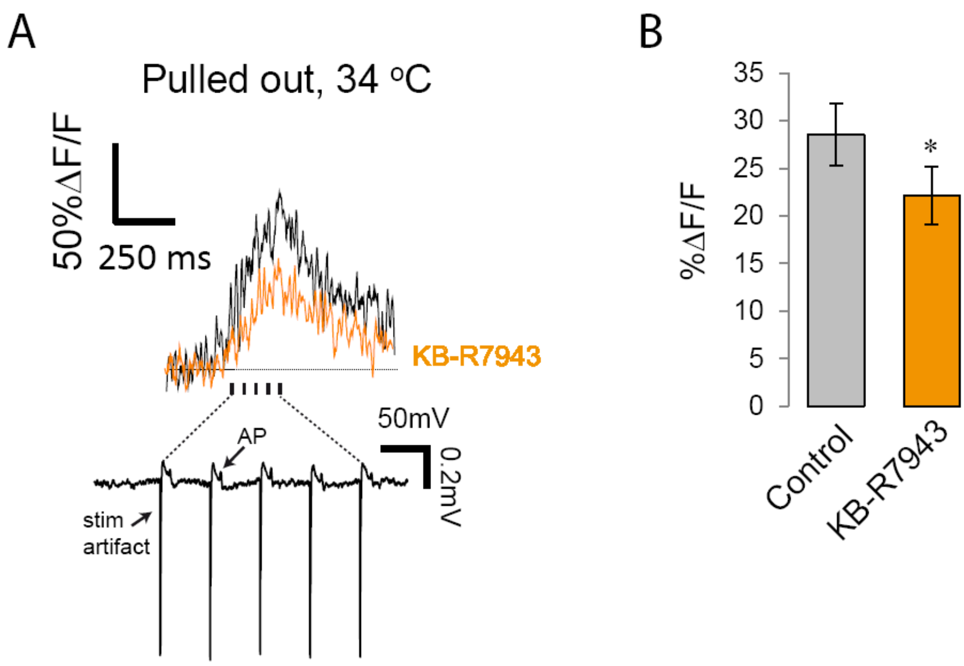


**Supplementary Figure 1. Effect of KB-R7943 on AIS spike-dependent $Ca^{2+}$ transients in non-clamped hippocampal granule cells from brain slices.**

(**a**) Spike-induced Ca2+ transients at the AIS of GCs loaded with Fluo-4. GCs were loaded with 400μM Fluo-4 and 80μM alexa594 for 2-3 minutes. After pulling-out the pipette cells were left to recover for 30 min at 34°C. Then, cells were re-patched in cell-attached mode, and axons were stimulated with 5 pulses at 20 Hz with a bipolar electrode inserted in the stratum lucidum (~500 μm far away from the soma). This protocol induced 5APs (recorded in cell-attached, lower trace in A) and similar $Ca^{2+}$ transients to those observed in whole-cell mode. (**b**) Summary of the effect of KB-R7943 in the amplitude of the transients.

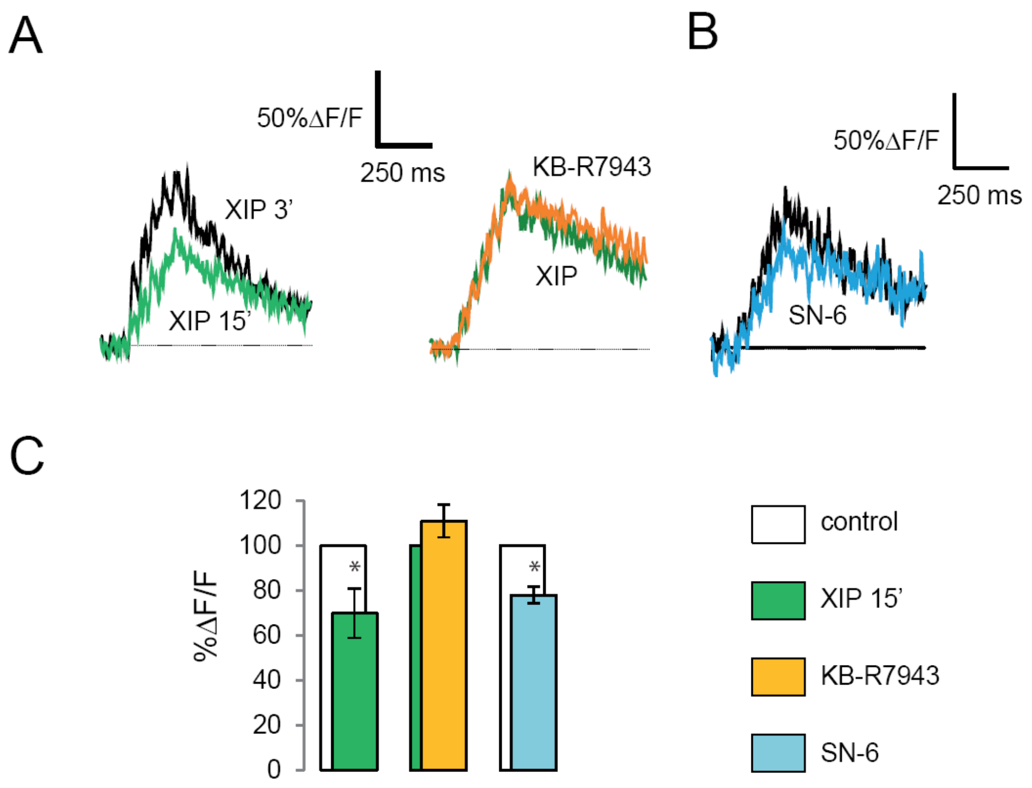


**Supplementary Figure 2. Effect of different NCX blockers on spike dependent $Ca^{2+}$ transients at the AIS of granule cells.**

(**a**) Example traces of the effect of XIP (*green trace*) on Ca2+ transients at the AIS 3 minutes after breaking-in. The stimulation protocol was 5 pulses at 20Hz. After 15 minutes in the presence of XIP (*green trace second panel*), KB-R7943 was applied to the bath (*orange trace*). XIP occluded the effect of KB-R7943. (**b**) Effect of SN-6 on Ca2+ transients at the AIS of GCs loaded with Fluo-4 for 10 minutes. (**c**) Summary of the results. Data are average of 6 cells per condition ±SEM. *p<0.05

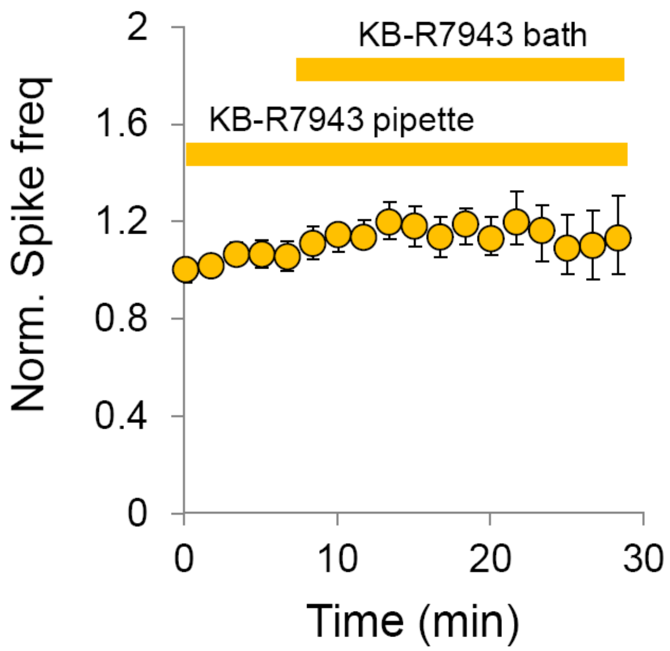


**Supplementary Figure 3. Testing possible indirect effects of KB-R7943 on single-cell spike frequency by affecting the network activity.**

Effect of bath-applied KB-R7943 on spike frequency in GCs loaded with KB-R7943 inside the pipette and with a high concentration of $Na^+$ (28 mM). Spikes were induced by a constant suprathreshold current pulse, as in experiments from figure 2.

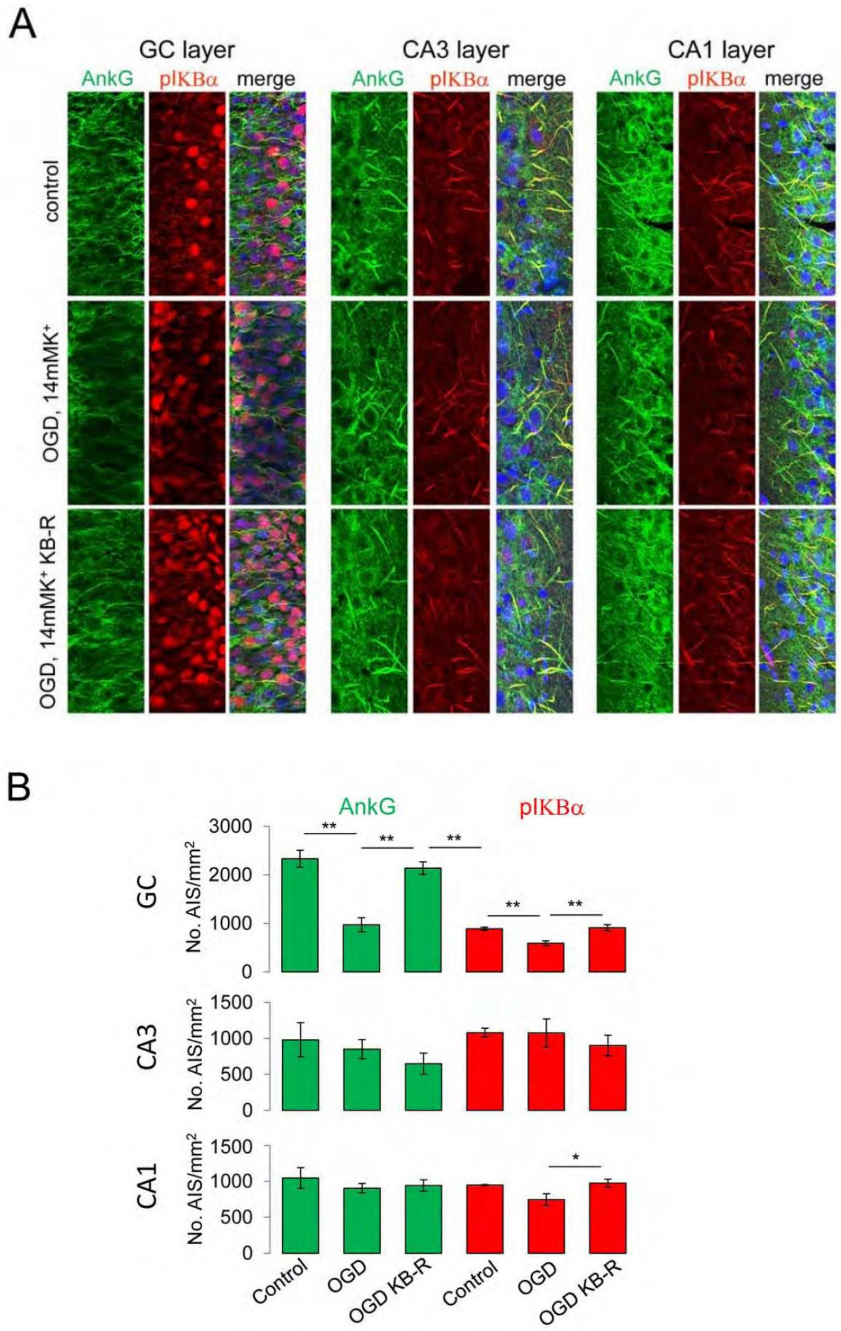


**Supplementary Figure 4**. **AISs of GCs, and not of CA3 and CA1 areas *in acute slices*, are dismantled after interruption of oxygen and glucose supply in high [$K^+$]. Blockade of $NCX_{rev}$ protects them.**

(**a**) Immunostaining for ankyrin-G and pIKBα in three different conditions: control, oxygen deprivation (No $O_2$ supply, no glucose, 14 mM $K^+$ for 1h) and OGD, 14mM$K^+$+KB-R7943 (2μM), as indicated in the figure. (**b**) Number of AISs for each experimental condition in GC, CA3 and CA1 areas. Data are mean ± SEM from 4 animals for each condition (5-6 slices/animal).

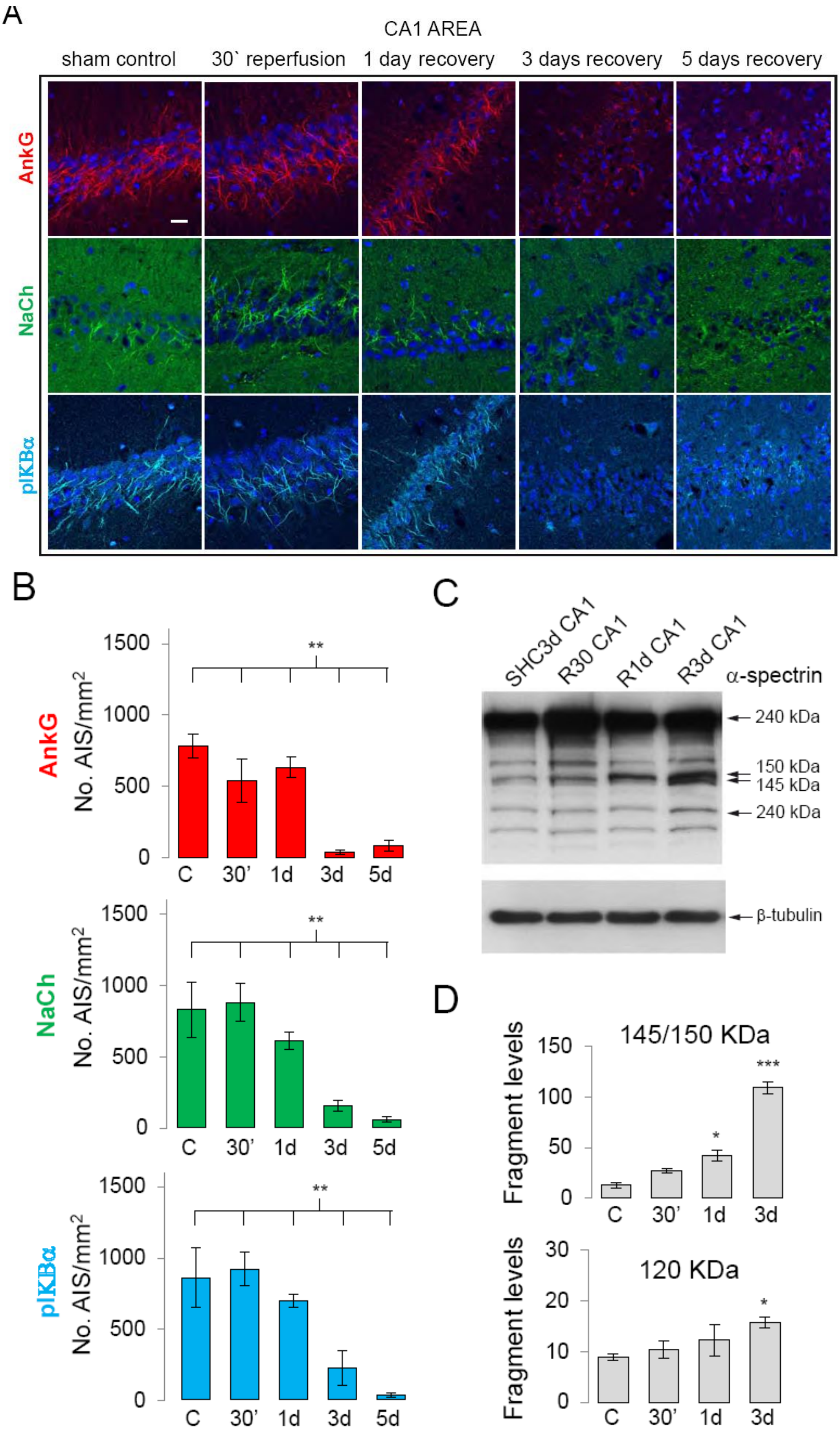
A
CA1 AREA
sham control
30` reperfusion
1 day recovery
3 days recovery
5 days recovery
AnkG
NaCh
pIKBα
B
No. AIS/mm2
1500
1000
500
0
**
C
30'
1d
3d
5d
C
SHC3d CA1
R30 CA1
R1d CA1
R3d CA1
α-spectrin
240 kDa
150 kDa
145 kDa
240 kDa
β-tubulin
D
145/150 KDa
Fragment levels
150
100
50
0
***
*
120 KDa
30
20
10
0
C
30'
1d
3d

**Supplementary Figure 5. Effect of ischemia on AISs dismantling in CA1 area *in vivo*.**

(**a**) Inmunostaining for Ankyrin-G, pIKB$\alpha$ and panNaCh in control pre-operated rats (sham), and in rats following variable periods of reperfusion: 30 min (30'), 1 day (1d), 3 days (3d) and 5 days (5d) following ischemia. (**b**) Summary of the results for the number of AIS in CA1 region for all periods of recovery. (**c**) Spectrin fragments in CA1 region after different recovery periods, as indicated (SHC3d corresponds to sham control at 3 days). (**d**) Average values for 145/150 kDa and 120KDa spectrin fragments (corresponding to calpain and caspase-3 activities, respectively). Values are averages of 4 rats for each condition ± SEM. Scale bar=30$\mu$m. *$p<0.05$; **$p<0.01$; ***$p<0.001$

A

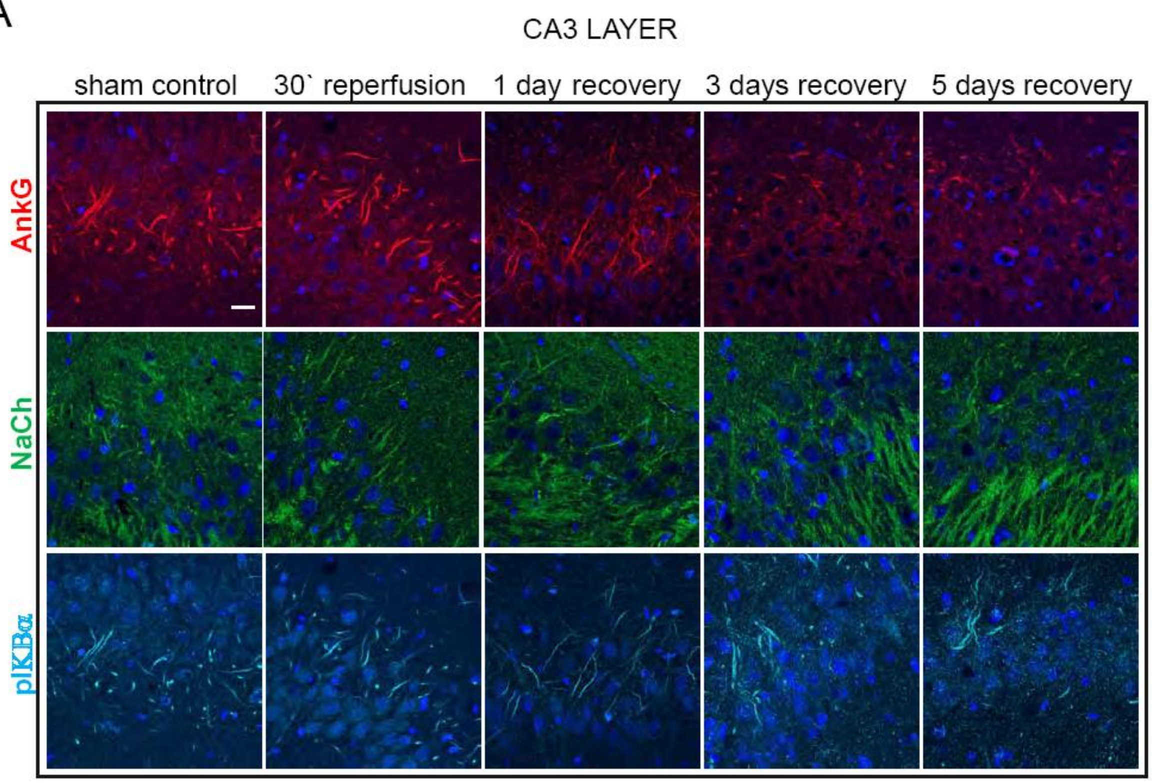


B

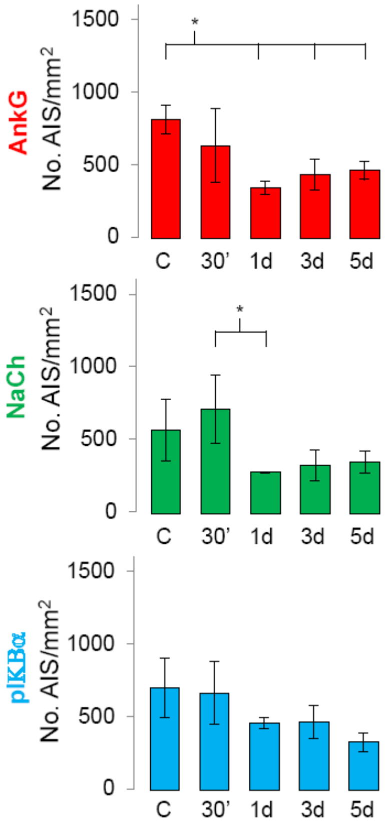


**Supplementary Figure 6. Effect of ischemia on AIS dismantling in CA3 area *in vivo*.**

(**a**) Inmunostaining for Ankyrin G, pIKBα and panNaCh in control pre-operated rats (sham) and in rats following variable periods of reperfusion: 30 min (30'), 1 day (1d), 3 days (3d) and 5 days (5d) after ischemia. (**b**) Summary of the results for the number of AIS in CA3 region for all periods of recovery. Values are averages of 4 rats for each condition ± SEM. Scale bar=30μm. *p<0.05; **p<0.01

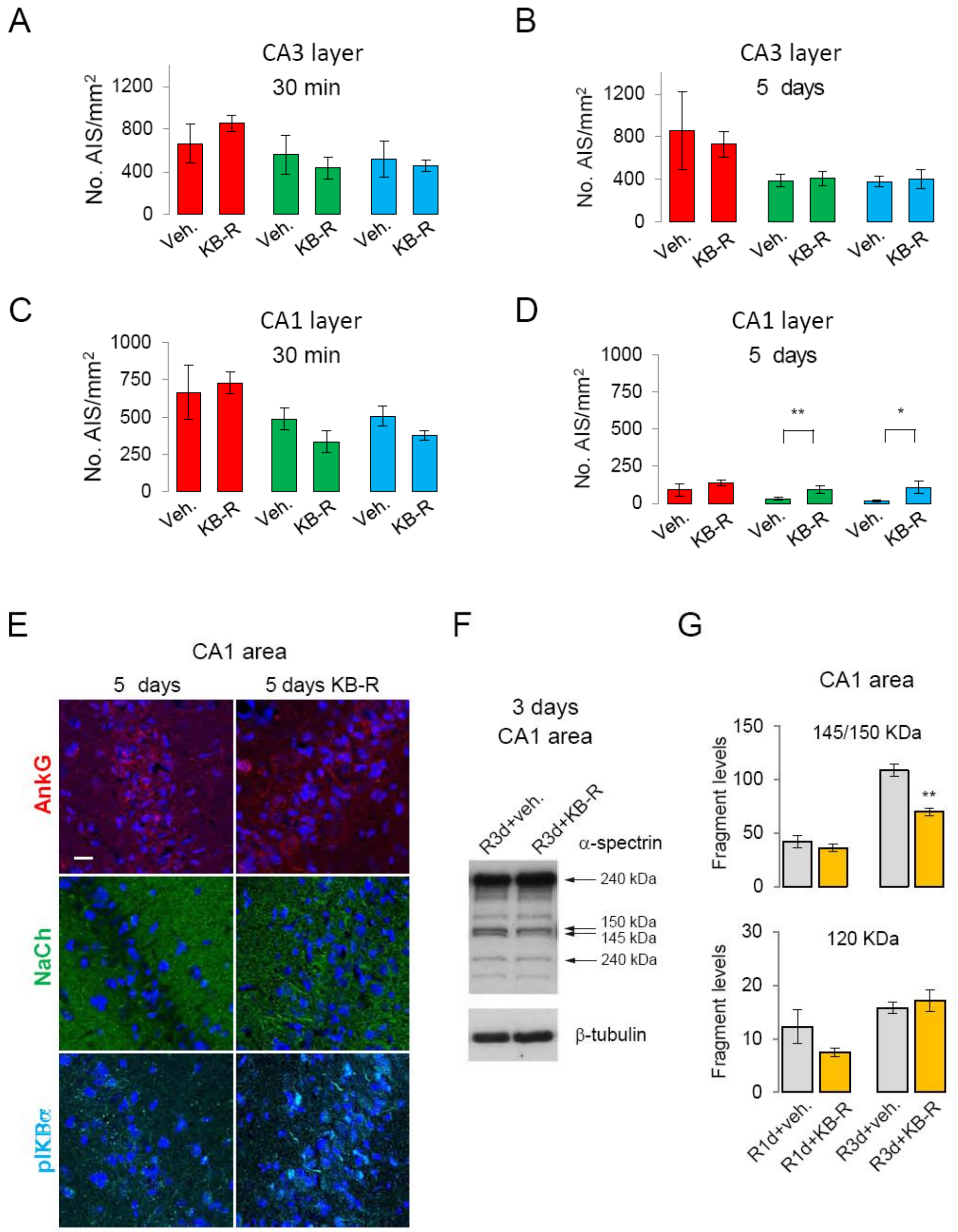


**Supplementary Figure 7. Effect of KB-R7943 on AISs dismantling in CA3 and CA1 areas after ischemia *in vivo*.**

(**a**) Average values of the number of AISs in vehicle or KB-R7943 treated animals in CA3 area at 30 min reperfusion after ischemia (*red*, ankyring G; *green*, Na+ channels; *blue*, pIKBα). (**b**) At 5 days of recovery after ischemia. (**c**) After 30 min reperfusion in CA1 area. (**d**) After 5 days of recovery in CA1 area. (**e**) Images corresponding to examples from panel D. (**f**) Spectrin fragments from CA1 area after 3 days from ischemia in rats treated with vehicle or KB-R7943. ***G***. Effect of KB-R7943 on spectrin fragment levels (145/150 kDa and 120 kDa, as indicated) after 1 day (R1d) or 3 days (R3d) following ischemia.

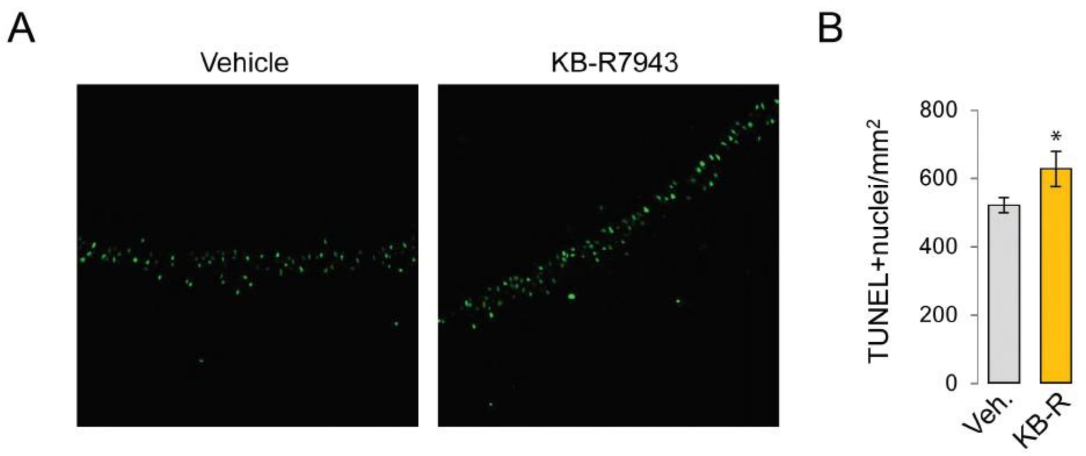


**Supplementary Figure 8. Effect of KB-R7943 on cell death in CA1 layer after 5 days of reperfusion following brief ischemia.** (**a**) TUNEL positive nuclei in CA1 area after 5 days of reperfusion in rats untreated (vehicle) or treated with KB-R7943. (**b**) Average ± SEM of 4 rats per condition. *p<0.05

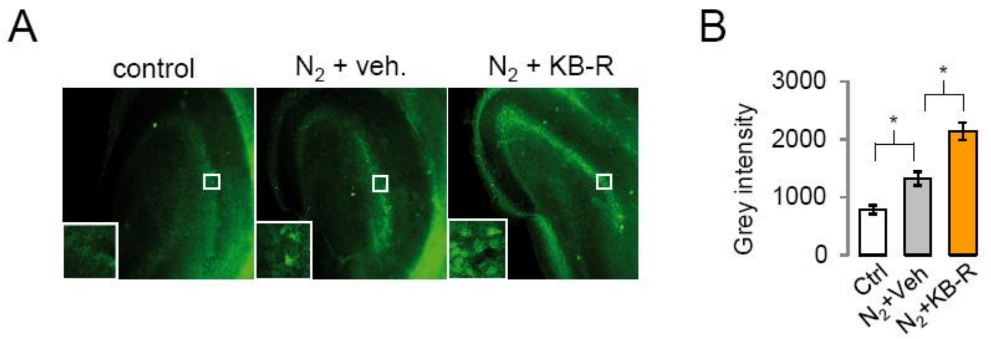


**Supplementary Figure 9. $Na^+/Ca^{2+}$ exchanger blockade increases cell damage in the dentate gyrus after oxygen deprivation in acute slices.**

(**a**) Example of fluorojade B staining of dentate gyrus after brief OGD in the presence or absence of KB-R7943. Insets show an enlargement of the granu cell layer. (**b**) Summary of the results. Data are mean ± SEM of 4 rats per condition.
